# Mineral dust stimulates microbial exoenzyme activity and enhances carbon mineralization capabilities in nutrient-poor peat soil

**DOI:** 10.64898/2026.07.28.741331

**Authors:** Jordan Thakar, E. Kathryn Hettinga, Kimber E. Munford, Susan Glasauer

## Abstract

Nutrient limitation is an important control on heterotrophic microbial activity that helps to stabilize the massive stocks of organic carbon held in ombrotrophic peatlands. Minerals contained in atmospheric dusts are critical nutrient sources for peatlands, yet the role of dust in supporting the below-ground microbial processes that underpin primary productivity is largely unknown. We investigated how mineral dust generated from mining waste rock (<20 µm) influences element bioavailability and subsurface microbial functioning using flow-through soil mesocosms. The bioavailability of base cations (Ca, Mg, K), transition metals (Fe, Al, Ni, Cu), Al, and P was tracked over two months at three soil depths (0-6, 6-12, and 12-18 cm) using an extended sequential extraction method. We also analyzed microbial community composition (16S rRNA and ITS amplicon sequencing) and mineralization capacity (exoenzyme assays and carbon substrate incubations). After two months, the concentration of metals in the peat increased substantially after dust application, but the mobility and bioavailability varied by element. Responses of microbial communities to dust application were highly dependent on depth from the surface. Carbon substrate incubations revealed enhanced mineralization capabilities in soil from the surface zone (0-6 cm), but a relatively low stimulation of exoenzymes. Soil pH and phosphorus mobility were also impacted near the site of dust application, while acid phosphatase activity was lower throughout the column. In the middle zone (6-12 cm), the activities of β-glucosidase, β-xylosidase, and NAGase were higher with dust exposure. Measured microbial activity mostly remained unchanged in the lowest depth (12-18 cm). We observed increases in the relative abundances of putative saprotrophic fungi throughout the mesocosm profile. Results from this experiment show that the deposition and weathering of mineral dust can induce a complex set of changes to the capacity and nature of microbial carbon mineralization within a shallow layer of peat.

## 1. Introduction

Peatlands encompass a diverse set of wetland ecosystems that are estimated to store up to one third of the Earth’s soil organic carbon (Yu *et al.*, 2010). Inputs of C into peat soils from plants and microorganisms are balanced against mineralization, in which microbes transform soil organic carbon into dissolved organic matter (DOM), carbon dioxide (CO_2_), or methane (CH_4_) (Limpens *et al.*, 2008). Land development and climate warming are recognized globally as severe threats to the preservation of peatland carbon (C) (Page *et al.*, 2022; Robinson *et al.*, 2023). Increases in the frequency of droughts can allow more oxygen to enter peat soils, which have been demonstrated to accelerate microbial C mineralization (Bu *et al.*, 2011; Barreto *et al.*, 2024). However, simultaneous boosts in primary productivity may offset carbon losses from the stimulation of microbial heterotrophs (Pinsonneault *et al.*, 2016; Hamard *et al.*, 2025). Besides temperature and soil moisture, factors such as poor substrate quality, limited nutrient supplies, and acidic pH regimes will also govern microbial community functioning in peatlands (Aerts *et al.*, 2001; Lin *et al.*, 2014).

Nutrient availability strongly limits microbial activity, especially in bogs, where fresh nutrient inputs are sourced solely from precipitation and atmospheric deposition (Bubier *et al.*, 2007). Mineralization of soil organic matter (SOM) by microbes is thus a critical mode of nutrient cycling that promotes ecosystem productivity. Microbes release extracellular enzymes (referred to as exoenzymes) to hydrolyze high molecular weight organic matter into monomers that can be imported into the cell as a source of carbon, nutrients, and energy (Allison and Vitousek, 2005). The energetic and nutritional costs of exoenzyme production and secretion make the hydrolysis of SOM a rate limiting step for microbial carbon and nutrient cycling in soils (Malik *et al.*, 2019). As a result, measuring the activity of exoenzymes can provide insights into the forms of organic matter that are targeted by microbial communities during different stages of mineralization (Mori *et al.*, 2021). Commonly measured exoenzymes include β-glucosidase and and β-xylosidase; these enzymes support carbon acquisition from glucosides in cellulose materials and xylans from hemicelluloses, respectively (Eivazi and Tabatabai, 1988; Campos *et al.*, 2014). Another is N-acetylglucosaminidase (NAGase), which contributes to N cycling from chitooligosaccharides in chitobiose or glycoproteins (Geisseler *et al.*, 2010). Acid phosphatase targets monophosphate esters in a broad suite of organic P compounds (Eivazi and Tabatabai, 1977).

Most investigations relating peatland productivity to nutrient availability tend to focus on the major macronutrients (nitrogen [N] and phosphorus [P]), with P typically the most limiting for ombrotrophic bogs (Myers *et al.*, 2012; Hill *et al.*, 2014; Pinsonneault *et al.*, 2016). Consequently, our understanding of how nutrient limitations shape C mineralization in peat does not consider metallic elements that could act as minor macronutrients (e.g., calcium [Ca], magnesium [Mg], and potassium [K]) or trace nutrients (e.g., iron [Fe], zinc [Zn], nickel [Ni], and copper [Cu]). Metallic nutrients have been demonstrated to influence microbial functioning in soil environments (Peng *et al.*, 2022; Dai *et al.*, 2023) and even improve estimates of soil respiration as a model input (Dhandapani *et al.*, 2021). Metals may serve as enzyme cofactors, structural components in cell membranes or walls, and as terminal electron acceptors (Warren *et al.*, 2017). Previous studies have investigated the impact of a narrow suite of metals on peatland microbial communities, either through fertilization experiments with simple mineral or dissolved amendments (e.g., Basiliko and Yavitt, 2001; Thomas and Pearce, 2004; Keller and Wade, 2018), or field observations along a nutrient or pollution gradient at a single timepoint (e.g., Luke *et al.*, 2015; Kujala *et al.*, 2018). These approaches have provided important insights into the sensitivity of peatland microbial communities to metal inputs but overlook a major pathway for nutrient additions to bogs: the deposition of multiple elements, simultaneously, by atmospheric dust. Dust hosts a wide range of potential metallic nutrient elements, depending on the source material (Arvin *et al.*, 2017). Current climate change is expected to increase the frequency and severity of dust events through intensifying storms and drought (Achakulwisut *et al.*, 2019; Huck *et al.*, 2023). Future mining developments, such as projects proposed for Ontario’s Ring of Fire, could increase atmospheric dust loads through road building and excavation. The generation of dust from the Athabasca tar sands (Alberta, Canada) and road development projects have already been correlated with detectable increases in the metal content of peat porewater and *Sphagnum* moss, along with influencing plant community structure in peatlands (Mullan-Boudreau *et al.*, 2017; Li *et al.*, 2023; Butt *et al.*, 2024). The long-term impacts of dust deposition on carbon sequestration in peatlands remains controversial (Kylander *et al.*, 2018; Schillereff *et al.*, 2021; Da Silva *et al.*, 2022). The direction and magnitude of the relationship between peatland carbon storage and dust will likely vary with dust composition and deposition rates. Incorporating the impacts of atmospheric mineral dust into our understanding of peatland carbon cycling will require a deeper understanding of how multiple elements simultaneously move through the soil environment and influence microbial functioning.

Our previous research has demonstrated that mineral dust can weather in acidic peat, release a suite of metals, and stimulate soil CO_2_ respiration within 1 to 2 weeks (Hettinga *et al.*, *in press*). In this study, we investigate the impacts of dust exposure on microbial community structure and activity as well as the bioavailability of metals in flow-through peatland mesocosms. We hypothesized that the mineral dust would stimulate microbial activity as dust particles break down in the acidic peat environment and release metallic. This stimulation in microbial activity was predicted to occur in tandem with shifts in microbial community structure that result in a larger representation of taxa with higher nutrient demands.

## 2. Materials and methods

### 2.1. Field site and peat collection

Peat soil was collected from Cranberry Bog, Puslinch, Ontario (43.465065, −80.218127) on October 27, 2022. The region surrounding the wetland is primarily agricultural, with several active and retired rock quarries within 100 km. The wetland complex consists of bog, fen, and marsh areas, with surrounding swamp lands. Samples were retrieved from the bog area; this area was identified by the acidic pH around 4.0 and low electrical conductivity of pore water collected in hollows (see Table S1), the accumulation of organic matter, and identifying bog vegetation such as *Sphagnum* spp. (green sphagnum moss), *Rhododendron* spp. (labrador tea), and *Andromeda polifolia* (bog rosemary) (CWCS, 1987). Peat blocks were excavated to a depth of 18 cm and immediately transported to the University of Guelph, where they were kept at 4℃ in the dark until the soil columns were prepared.

### 2.2. Mineral dust collection and characterization

Fine mineral sediment was collected from a waste rock pile at a legacy nickel-copper mine site managed by the Ontario Ministry of Energy, Northern Development and Mines near Sudbury, Ontario in July of 2020. The mine was operational until 2015. The waste rock was not acid generating and consisted primarily of feldspar, quartz and clay minerals (Munford, 2024). Ground rock was sieved to <20 µm using a Gilson GilSonic UltraSiever Sonic Sifter GA-8. A broad elemental characterization of the rock material was performed on the <20 µm fraction size with a reverse aqua regia digestion (USEPA, 2007) with the inclusion of the EnviroMAT SS-2 Soil Standard (SCP Science) as the reference material. Elements of interest were then analyzed with Inductively Coupled Plasma-Optical Emission Spectroscopy (ICP-OES) (see Table S2).

### 2.3. Flow-through peatland mesocosms

The living plant layer was removed from the peat blocks before preparing cores, in order to separate the most biologically active zone for heterotrophic soil microbes (*i.e.,* the acrotelm) (see Figure S1). Cores with a diameter of 10 cm and a depth of 18 cm were cut from the blocks and placed into translucent PVC tubes (10 cm x 30 cm). The cap at the base of the tube contained an outlet to allow for drainage of pore water. The top of each peat mesocosm was loosely covered with fine nylon mesh to inhibit dust contamination between cores while also maintaining the connection to the ambient atmosphere (see Figure S2).

To treat the peat soil with the mineral dust (treatment PD), the prepared dust was applied to the top surface of the mesocosms, while the control treatments did not receive the dust amendment (control P). Dust was added to the treated columns based on the total dry weight of the top 6 cm of each peat core using a ratio of 0.127 g dry dust to 1 g dry peat. This ratio was chosen to match the ratio used in a related study (Hettinga *et al.*, *in press*). The total amount of dust added to each core ranged between 2.56 and 3.40 g dry dust. Dust was applied evenly across the top of the cores through fine nylon mesh, resulting in surface concentrations ranging between 3.2 x 10^-2^ and 4.3 x 10^-2^ g dry dust · cm^-2^. Each column was incubated in the dark at 23℃ and received weekly inputs of sterile simulated rainwater (Anderson *et al.*, 2000). The water level of each column was maintained near 8 cm from the top, with approximately 50 mL of porewater replaced weekly. The mesocosms were incubated for two months with sacrificial sampling in triplicate at the beginning of the experiment (T0), one month (T1), and two months (T2), for a total of 15 columns. Select peat columns were removed at each timepoint and sectioned by respective depth: 0-6 cm (D1), 6-12 cm (D2), and 12-18 cm (D3). These depth categories encompassed the upper zone with unsaturated pores, the partially saturated zone, and the consistently saturated layer, respectively.

To avoid edge effects, column sections were first halved by length after removal from the acrylic sleeve and sampled from the center of the column for DNA extraction and sequencing; these samples were stored at −20℃ until processing. The remaining peat was homogenized at each respective depth, then divided into portions to assess extracellular enzyme activity, carbon mineralization capacity, and soil chemical parameters. Exoenzyme and carbon mineralization measurements were performed within 3 weeks after storage at 4°C. For the sequential metal extractions, peat samples were lyophilized, ball milled, and stored at −20℃ until extraction. Weekly pore water samples were filtered (0.45 µm), acidified (2% HNO_3_) and analyzed for dissolved inorganic elements by ICP-OES.

### 2.4. Exoenzyme activity assays

The potential activities of acid phosphatase, β-glucosidase, β-xylosidase, and N-acetylglucosaminidase (NAGase) were measured using standard microplate colorimetric assays (McKergow *et al.*, 2021). For a given exoenzyme, soil samples were suspended in a pH-buffered solution and incubated with a target substrate. The target substrate is tagged with a fluorophore that is activated upon cleavage by the exoenzyme. Incubation times and concentrations used for target substrates are provided in Table 1. Five sample and five analytical replicates were performed for each sample and enzyme. In each well, 200 µL of soil suspension was combined with 200 µL of target substrate. Absorbance values were measured at 405 µm, using a Biotek ELx800 Microplate Reader. Measured absorbances were converted to potential enzyme activity (µmol · h^-1^ · g soil^-1^) with a PNP standard curve (Center for Dead Plant Studies 1994; McKergow *et al.*, 2021).

**Table 1:**
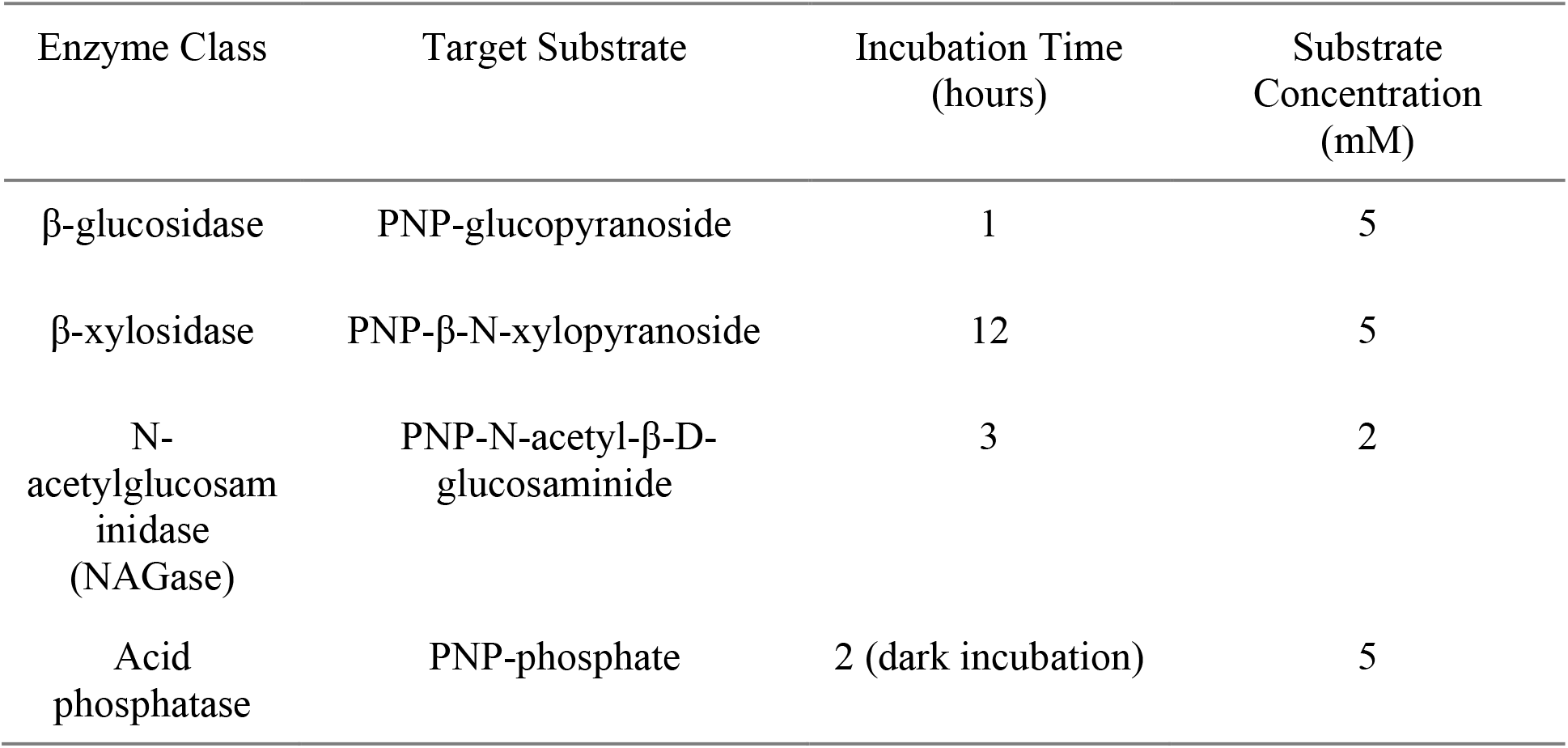
General summary of colorimetric assays used for measuring target exoenzymes.

### 2.5. Community Level Physiological Profiling

Community level physiological profiling (CLPP) was conducted using Biolog Ecoplates^TM^. Soil suspensions for analysis were made by vortexing 7 mM phosphate-buffered saline solution (PBS) at pH 7 with peat samples at a ratio of 4 g: 40 mL peat to PBS. One Biolog Ecoplate^TM^, containing 31 carbon sources in triplicate, was inoculated for each sample using 100 µl of soil-buffer suspension, then wrapped in aluminum foil to block light, and placed on a shaker at 75 rpm. Absorbance values were measured at 510 µm wavelength every 12 hours for 7 days (Biotek ELx800 Microplate Reader). Average absorbance values for each carbon substrate are reported as sums by the appropriate functional group classification including amines, amino acids, carbohydrates, carboxylic acids, phenolics, and polymers (Sala *et al.*, 2010).

### 2.6. DNA extractions and microbiome analysis

DNA extractions of peat samples were completed using the Metagenom Bio Soil DNA Extraction Kit according to the manufacturer’s protocols (Metagenom Bio Life Science Inc, Waterloo, ON), with the additional step suggested by the manufacturer to remove inhibitory humic molecules. The quality and quantity of DNA in extracts were measured with a Nanodrop 8000 Spectrophotometer, and DNA quantity was further verified with Qubit fluorimetry. Five extraction replicates were used for each sample, and replicates were pooled prior to being submitted to Metagenom Bio Life Sciences (Waterloo, Ontario, Canada) for amplicon sequencing on an Illumina MiSeq. For the prokaryotic 16S rRNA gene target, DNA extracts were amplified using universal primers 515FB (5’-GTGYCAGCMGCCGCGGTAA) and 806RB (5’-GGACTACNVGGGTWTCTAAT) (Walters *et al.*, 2015). The primers BITS (5’-ACCTGCGGARGGATCA) and B58S3 (5’-GAGATCCRTTGYTRAAAGTT) were used for targeting the fungal ITS gene (Bokulich and Mills, 2013). Primers were removed from demultiplexed Illumina 16S rRNA and ITS gene sequences with QIIME2 (Bolyen *et al.*, 2019) and forwarded to R for processing through the DADA2 pipeline (version 1.26) (Callahan *et al.*, 2016). We aligned Amplicon Sequencing Variants (ASVs) to the SILVA database (release 138) and UNITE database (version 10) to assign taxonomy to 16S rRNA and ITS reads, respectively (Quast *et al.*, 2013; Abarenkov *et al.*, 2024). Any ASVs classified as chloroplast and mitochondria ASVs were removed. After processing, 9055 bacterial ASVs and 2678 fungal ASVs remained. Sequences were compiled into phyloseq objects for further analysis using the *phyloseq* R package (McMurdie and Holmes, 2013). The following alpha diversity metrics were computed from ASV counts using the *phyloseq* R package: observed ASV richness (R), Berger-Parker Index (BP), and Shannon Diversity Index (H’). Each of these alpha diversity metrics were chosen to characterize different contributors to alpha diversity based on the guidelines provided by Cassol *et al.* (2025). Richness is defined as the total count ASVs in each sequencing library. The Berger-Parker Index was used to assess the distribution of relative abundances between ASVs in a sample (referred to as community “evenness”). Shannon Diversity is a metric that reflects changes in richness and community evenness. Rarefaction curve analysis indicated that all amplicon samples for the 16S rRNA gene were sequenced with enough depth to accurately compare alpha diversity metrics (see Figures S3-S4). One sample that was sequenced for ITS amplicons did not have a saturated rarefaction curve and this sample was omitted when comparing alpha diversity. Amplicon samples were not rarefied for any downstream analyses. To account for compositionality inherent in marker gene sequencing, all ASV data was transformed with centered-log ratio (clr) prior to ordinations, permutational tests, and differential abundance analysis (Gloor *et al.*, 2017). To test for significant differences in microbial community composition and variance within sample groups, we performed permutational multivariate analysis of variance (PERMANOVA) and permutational dispersion analysis (PERMDISP) using an Aitchison distance matrix with the *vegan* R package (Oksanen *et al.*, 2025). Differential abundance analysis was conducted with the *ALDEx2* R Package (Fernandes *et al.*, 2014) to identify whether bacterial and fungal ASVs were significantly different in abundance between the dust treatment and control. Exploration of community structures with ordinations and compositional bar plots were generated using the *microViz* R package (Barnett *et al.*, 2021). Putative trophic modes and ecological guilds were assigned to fungal ASVs with the FUNGuild database (version 1.1) (Nguyen *et al.*, 2016).

### 2.7. Soil chemistry: soil pH, sequential extraction and reactive phosphorus analysis

Soil pH was measured in duplicate using the CaCl_2_ method (Ziadi and Tran Sen, 2007). An extended sequential extraction protocol was used to characterize the abundance and distribution of base cations, trace metals, and phosphorus across different soil fractions in the peat. The sequential extraction protocol used is a modified version of the procedure described by Leermakers *et al.* (2019), with the addition of a water-soluble fraction to assess element concentrations in a highly bioavailable form (Table 2) (Hass and Fine, 2010). Five sample replicates were performed per depth at T0 and T2 months; extracts were then filtered (0.22 µm), acidified (2% HNO_3_), and stored at 4°C. The extracts were analyzed for K, Ca, Mg, Fe, Al, Ni, Cu, Zn, and P using ICP-OES (Varian Vista Pro CCD Simultaneous ICP-OES). The Montana II soil standard was used as the reference standard (Millipore Sigma, NIST2711A).

**Table 2:** General summary of the sequential extraction procedure.

| Fraction Number | Fraction | Reagent | Duration of Extraction | Extraction Mechanism |
| --- | --- | --- | --- | --- |
| F1 | Water-soluble | MilliQ water | 16 hours | Dissolution |
| F2 | Exchangeable | 1 M Ammonium | 16 hours | Cation |
|  |  | acetate buffer<br>(pH 5) |  | exchange and<br>complexation |
| F3 | Labile organics | 0.1 M Sodium<br>pyrophosphate<br>($\text{Na}_4\text{P}_2\text{O}_7$) | 2 rounds: 1 hour<br>each | Complexation or chelation<br>of labile<br>organics (e.g.<br>humic and<br>fulvic acids) |
| F4 | Amorphous oxides and<br>hydroxides (principally<br>iron and aluminum) | 1.7 M Oxalate<br>buffer<br>($\text{NH}_4$)$_2$ $\text{C}_2\text{O}_4$ +<br>$\text{H}_2\text{C}_2\text{O}_4$ (pH 3.8) | 6 hours | Weak<br>reduction |
| F5 | Residual (includes<br>refractory organics,<br>crystalline oxides,<br>sulfides, and silicates) | 16 M Nitric Acid<br>($\text{HNO}_3$), and 12<br>M hydrochloric<br>acid (HCl) | 48-hour cool<br>digestion and 16-hour hot digestion | Mineral<br>dissolution |

### 2.8. Visualization and statistical analysis of chemical and microbial activity data

All data analysis and visualization were performed using the R programming language (R version 4.5.1). Most statistical tests were performed using the *stats* R package (R Core Team, 2025). Shapiro-Wilk tests revealed that all data were not normal (see Table S5). Kruskal-Wallis (KW) tests were used to compare mean values among different sample groups. If a KW test was significant, pairwise comparisons of mean values were then conducted with a post-hoc Dunn’s test using the *FSA* R package (Ogle *et al.*, 2025). Pairwise comparisons between the treatment and control (PD and P) for a given biological or chemical dataset are outlined in the main text; however, all pairwise comparisons between depths, timepoints, treatments, or combinations of these variables are reported in Tables S6-S10. The high degree of collinearity amongst the tested chemical variables prevented us from performing any meaningful correlation tests between chemical and biological datasets. Instead, we propose biologically relevant relationships between biological activity, peat chemistry, and the dust treatment in the discussion section.

## 3. Results

### 3.1. Exoenzyme activity

The activities of soil exoenzymes were measured to characterize broad shifts in nutritional demands by microbial communities based on the degradation of extracellular organic compounds. The measured activities for all tested exoenzymes had generally declined by T1 in both control peat (P) and peat treated with dust (PD) (Figure 1). Declining activity at T1 was more severe for the PD treatment compared to the control. This reduction in activity was most pronounced in the upper zone, but β-glucosidase activity was significantly lower than the control throughout the core (Z_PD-P_ = 4.47, p < 0.001).

**Figure 1.**
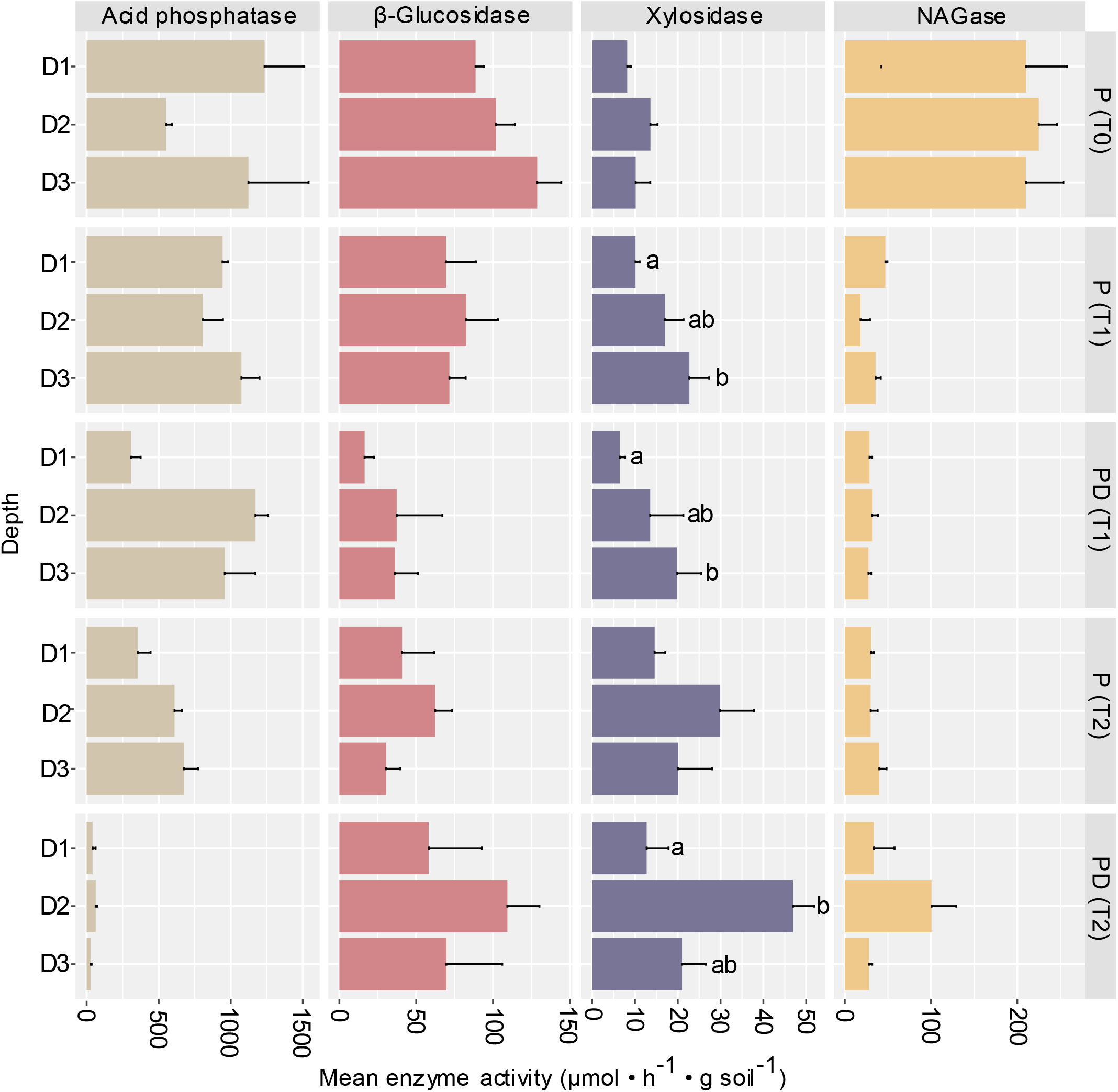
Mean activity (µmol · h^-1^ · g soil^-1^) of hydrolytic exoenzymes including acid phosphatase, β-glucosidase, xylosidase, and N-acetylglucosaminidase (NAGase). Samples are organized by treatment (peat with dust [PD] and peat without dust [P]), timepoint (T0, T1, and T2 months), and peat depth (D1: 0-6, D2: 6-12, and D3: 12-18 cm). Letters indicate significant differences in mean enzyme activity between peat depths within the same treatment and timepoint (*i.e.,* a significant result from Kruskal-Wallis and Dunn’s post-hoc test).

The activity of all exoenzymes, apart from acid phosphatase, increased between T1 and T2 in the PD treatment. This measured stimulation of exoenzyme activity for the dust treatment was highest in the 6-12 cm depth from the top (D2), where the activities of β-xylosidase, β-glucosidase, and NAGase were 57%, 75%, and 195% higher than the control, respectively. By T2, the activities of β-xylosidase, β-glucosidase, and NAGase in depths 0-6 cm (D1) and 12-18 cm (D3) of the PD treatment were similar to the control. Acid phosphatase was the only tested exoenzyme to significantly decline in the PD treatment after T1 (Z_T2-T1_= 3.18, p < 0.001), ranging between 3-10% of the corresponding controls by T2. When averaged across the column, acid phosphatase activity was significantly lower in the PD treatment compared to the control (Z_PD-P_ = 5.12, p < 0.001).

### 3.2. Growth experiments on Biolog Ecoplates^TM^

We used Biolog Ecoplates^TM^ to investigate how the dust treatment may alter microbial utilization of 31 different carbon sources through time and across different peat depths. Total microbial carbon mineralization was estimated by calculating the average absorbance for a substrate across all replicates and taking the sum of all 31 substrates for a given sample group (Figure 2). Similar to the activity measurements generated with exoenzyme assays, the total degradation of substrates on Ecoplates^TM^ had significantly declined by T1 in both the control and PD treatment relative to T0 (p < 0.01). Declines in substrate utilization between T0 and T1 were larger for the PD treatment in D1 and D2 relative to the control.

**Figure 2.**
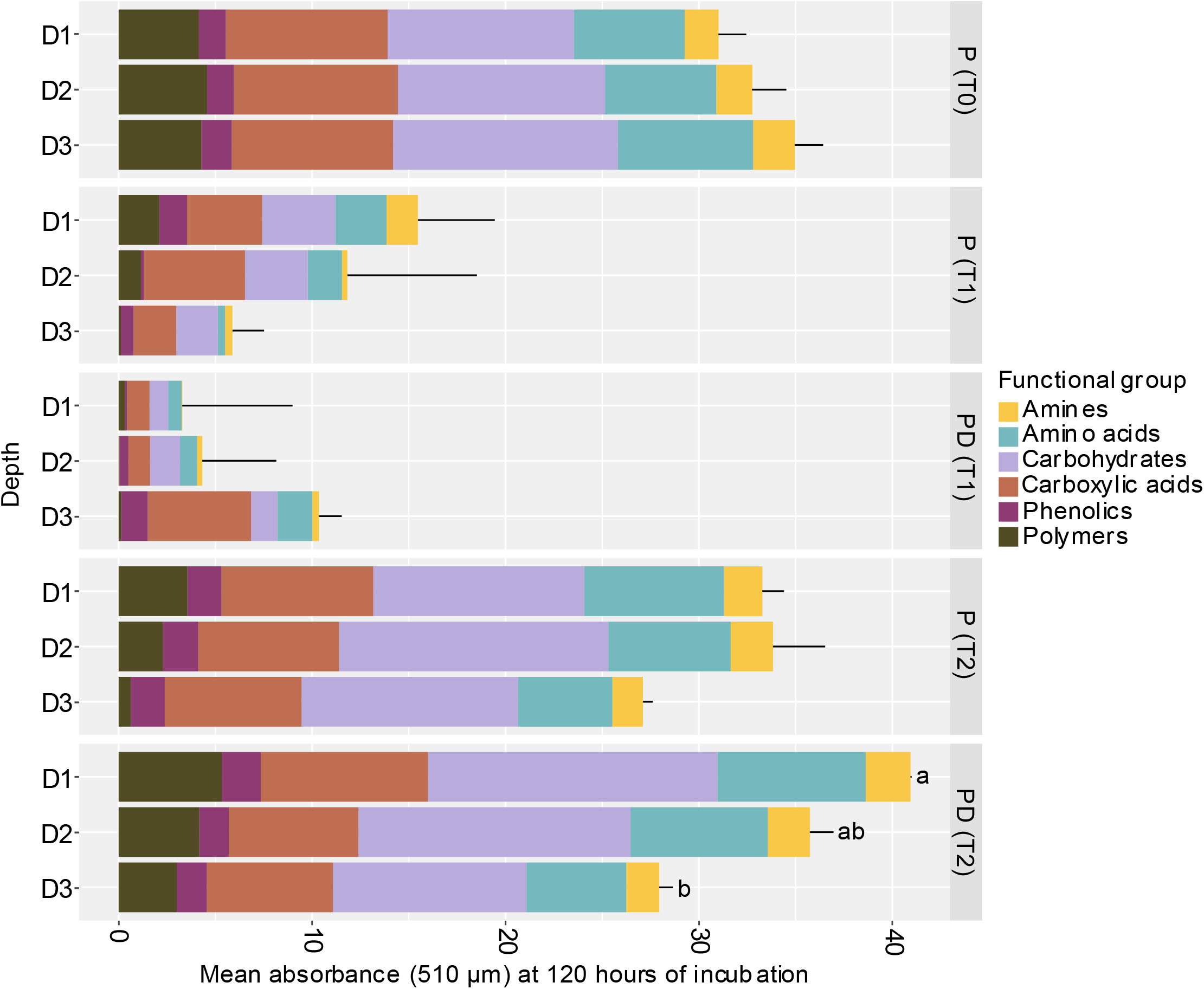
Sum of mean absorbance values (510 µm) for each carbon functional group after 120 hours of incubation on Biolog Ecoplates^TM^. Absorbance is proportional to the degree of carbon mineralization for the 31 unique carbon substrates, which are organized by carbon functional group (amines, amino acids, carbohydrates, carboxylic acids, phenolics, and polymers). Samples are organized by treatment (peat with dust [PD] and peat without dust [P]), timepoint (T0, T1, and T2 months), and peat depth from the top (D1: 0-6, D2: 6-12, and D3: 12-18 cm). Error bars represent the variation of the summed absorbance value for a sample across three replicate incubations on the same plate. Letters indicate significant differences in the total absorbance value (summed across functional groups) between peat depths within the same treatment and timepoint (*i.e.,* a significant result from Kruskal-Wallis and Dunn’s post-hoc test).

By the end of the experiment at T2 months, total carbon mineralization had significantly increased in both the PD treatment and control relative to T1 (P: Z_T2-T1_= 2.84, p = 0.04; PD: Z_T2-T1_= 4.43, p < 0.001). Substrate mineralization was highest in D1 of the PD treatment at T2 compared to the control. The mean measure of total carbon mineralization for the PD treatment at D1 was 6.8% higher than the control (P: 33.3 ± 1.1; PD: 40.1 ± 0.1), and 8.9% higher than the control at T0 (31.01 ± 1.44). Substrate mineralization capacity in D2 and D3 were more similar between the treatment and control by the end of the experiment. Comparisons of substrate mineralization between timepoints and treatments were consistent when assessing each carbon group individually (*i.e.,* amines, amino acids, carbohydrates, carboxylic acids, phenolics, and polymers). Out of all the functional groupings, mineralization capacity of polymerized substrates was enhanced the most in D1 of the PD treatment (P: 2.02 ± 0.03; PD: 4.16 ± 0.5), followed by carbohydrates for the same depth and treatment (P: 7.67 ± 0.78; PD: 14.96 ± 0.25) (Figure 2).

### 3.3. Prokaryotic and fungal community structures

We used amplicon sequencing of the 16S rRNA and ITS genes to monitor the structure of prokaryotic and fungal communities across all treatments, timepoints, and depths. Distinctions in the taxonomic structure of microbial communities between the PD treatment and control were prevalent throughout the mesocosm but depended on the depth of the peat. Prokaryotic community structures were dominated by bacteria; the relative abundance of archaea was <1% in all samples. We identified 48 prokaryotic phyla and classified most ASVs to the family level (n = 228 families). Acidobacteriota and Proteobacteria were the dominant phyla present in all samples, followed by Verrucomicrobiota, Bacteroidota, Myxococcota, and Desulfobacterota. With respect to fungi, 12 phyla were identified and most ASVs were classified to the genus level (n = 247 genera). Ascomycota and Basidiomycota were the major phyla identified, followed by Mortierellomycota, Rozellomycota, and ASVs only identified to the Fungi kingdom.

Fungal richness in both PD and control samples tended to decrease at T1 and then increase by T2 (Table S12). By the end of the experiment, fungal richness in the PD treatment was lower than the control in D1 (P_D1_: R = 422; PD_D1_: R = 352) and D2 (P_D2_: R = 560; PD_D2_: R = 356), but higher in D3 (P_D3_: R = 388; PD_D3_: R = 437). Prokaryote richness tended to decline throughout the experiment in both the PD treatment and control at all depths (see Table S11). By T2 months, the PD treatment had lower prokaryote richness than the control at D1 (P_D1_: R = 650; PD_D1_: R = 478) and D2 (P_D2_: R = 958; PD_D2_: R = 862), but a similar value at D3. Community evenness was estimated with the Berger-Parker Index (BPI). Fungal community evenness was higher in all depths of the PD treatment when compared to the control at T2, especially in D3 (P_D3_: BPI = 0.24; PD_D3_: BPI = 0.15). Prokaryotic community evenness was less impacted by the dust treatment, with BPI values ranging between 0.01 to 0.035 for both PD and control samples. The structures of the fungal and prokaryotic communities were not significantly different between the PD treatment and control according to PERMANOVA tests (Table 4). However, we still observed differences in the relative abundances of several taxa between the PD treatment and control at T1 and T2 months. Changes to fungal community structure were unique to each depth, but often involved genera belonging to the classes Agaricomycetes, Geoglossomycetes, and Saccharomycetes (Figure 4). After 1 month, distinctions between the PD treatment and control were driven by genera *Archaeorhizomyces* in the middle depth (P = 0.12%; PD = 36.33%), and *Phialocephala* in the lowest depth (P = 4.01; PD = 35.96%). By the end of the experiment at T2, the relative abundance of the *Archaeorhizomyces* genus was higher in the PD treatment (PD = 4.08%; P = 0.17%). Other taxonomic shifts in D1 at T2 occurred in ASVs that could only be identified to groups above the genus level, included ones classified to the Fungi kingdom (P =7.03%; PD = 14.89%) and Ascomycota phylum (P = 1.72%; PD = 3.90%), along with decreases for those classified to the Helotiales order (P = 19.35%; PD = 12.77%) and Basidiomycota phylum (P = 9.31%; PD = 0.73%). In the middle depth, the relative abundances of the following genera were higher in the PD treatment after 2 months: *Tomentella* (P = 0.62%; PD = 10.26%), *Hypholoma* (P = 4.06%; PD = 16.93%), *Hyaloscypha* (P = 7.04%; PD = 13.49%), and *Scopuloides* (P = 0.30%; PD = 5.46%). The relative abundances of several fungal genera also shifted in the lowest depth with the PD treatment after 2 months, such as increases in *Serendipita* (P = 0.97%; PD = 13.48%), *Trichoglossum* (P = 0.26%; PD = 15.54%), and *Hygrocybe* (P = 2.66%; PD = 11.92%), but a decrease in *Cyberlindnera* (P = 21.54%; PD = 0.40%).

Ecological functional groups were predicted for fungal ASVs using the FUNGuild database, which includes putative saprotrophs, symbiotrophs, pathotrophs, or some combination of the three. A majority of annotated ASVs were putative saprotrophs, followed by putative symbiotrophs (see Table S14 and Figure S5). The PD treatment had a higher average relative abundance of putative saprotrophs across all depths at T2 months (P = 42.3% ± 0.9; PD = 35% ± 2.5). A higher relative abundance of putative saprotrophs with the PD treatment was most evident in D2 (P = 12.4%; PD = 38.4%), and D1 to a lesser extent (P = 27.6%; PD = 36.3%).

Compositional increases in putative saprotrophs were mostly associated with three genera including *Hyaloscypha*, *Hypholoma*, and *Hygrocybe*. Approximately 37% of fungal ASVs were assigned to a putative guild, representing an average relative abundance of 58% ± 0.11 across all samples.

Differences in prokaryotic community structures between the PD treatment and control were frequently related to the Acidobacteriae class (Figure 3), but these differences in relative abundances were smaller than observed for fungi. After two months, the relative abundance for several prokaryotic taxa was higher in D1 of the PD treatment compared to the control, including ASVs identified to the Acidobacteriaceae (Subgroup 1) family (P = 5.39%; PD = 9.20%), the Koribacteraceae family (P = 5.29%; PD = 8.41%), and the Acidobacteriales order (P = 8.32%; PD = 11.35%). In contrast, the relative abundances of ASVs identified to these same taxonomic groups were lower in D3 of the PD treatment compared to the corresponding control. Prokaryotic community structures in D2 were similar between the PD treatment and the control at T2 (Figure 3), with differences in the relative abundance for the most dominant prokaryotic taxa remaining ≤1% (Figure 3).

**Figure 3.**
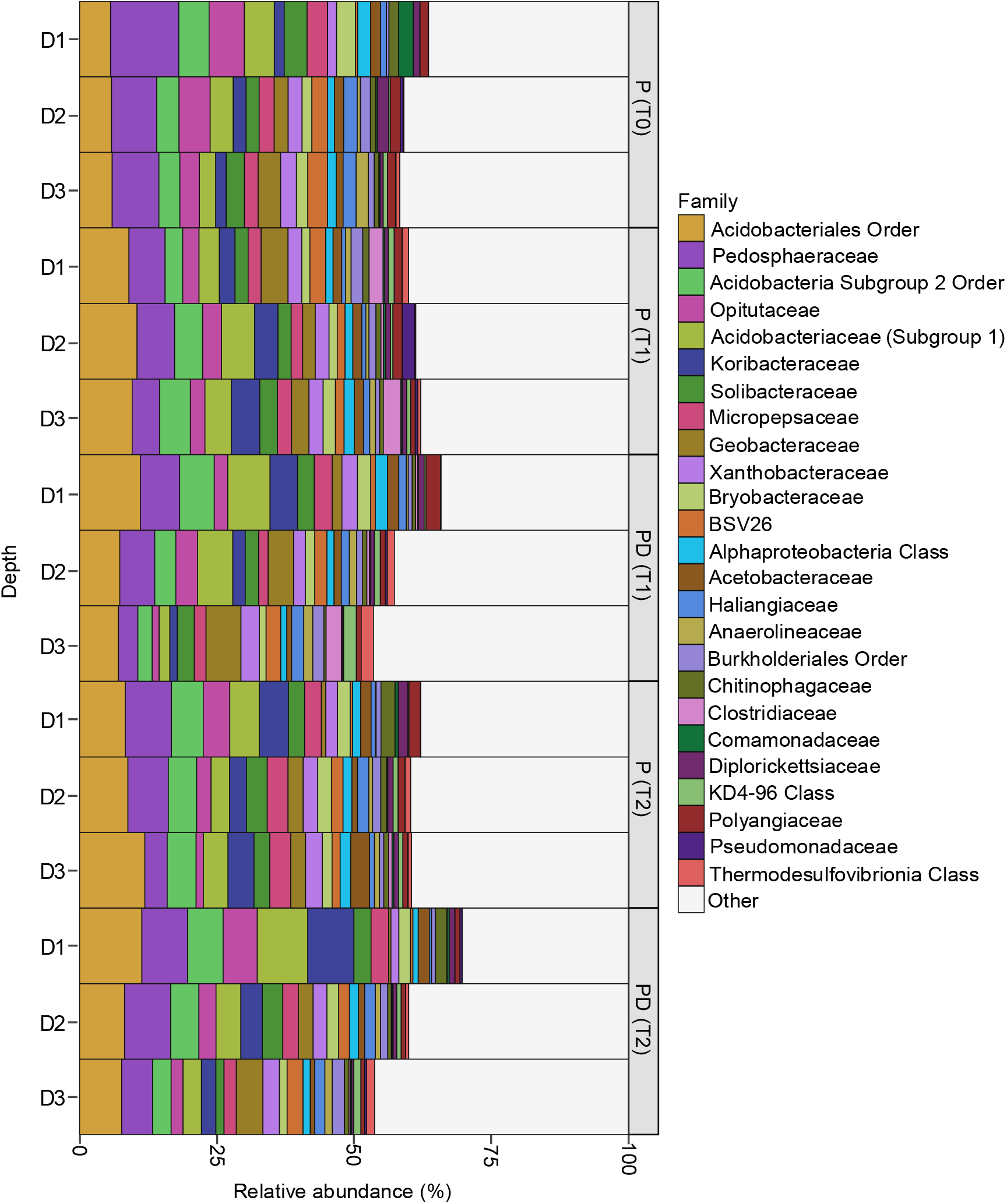
Relative abundance of prokaryotic families identified with 16S rRNA gene sequencing. Samples are organized by treatment (peat with dust [PD] and peat without dust [P]), timepoint (T0, T1, and T2 months), and peat depth (D1: 0-6, D2: 6-12, and D3: 12-18 cm). Prokaryotic families displayed in the legend were prevalent in at least one sample and had a total relative abundance >2% when summed across all samples.

**Figure 4.**
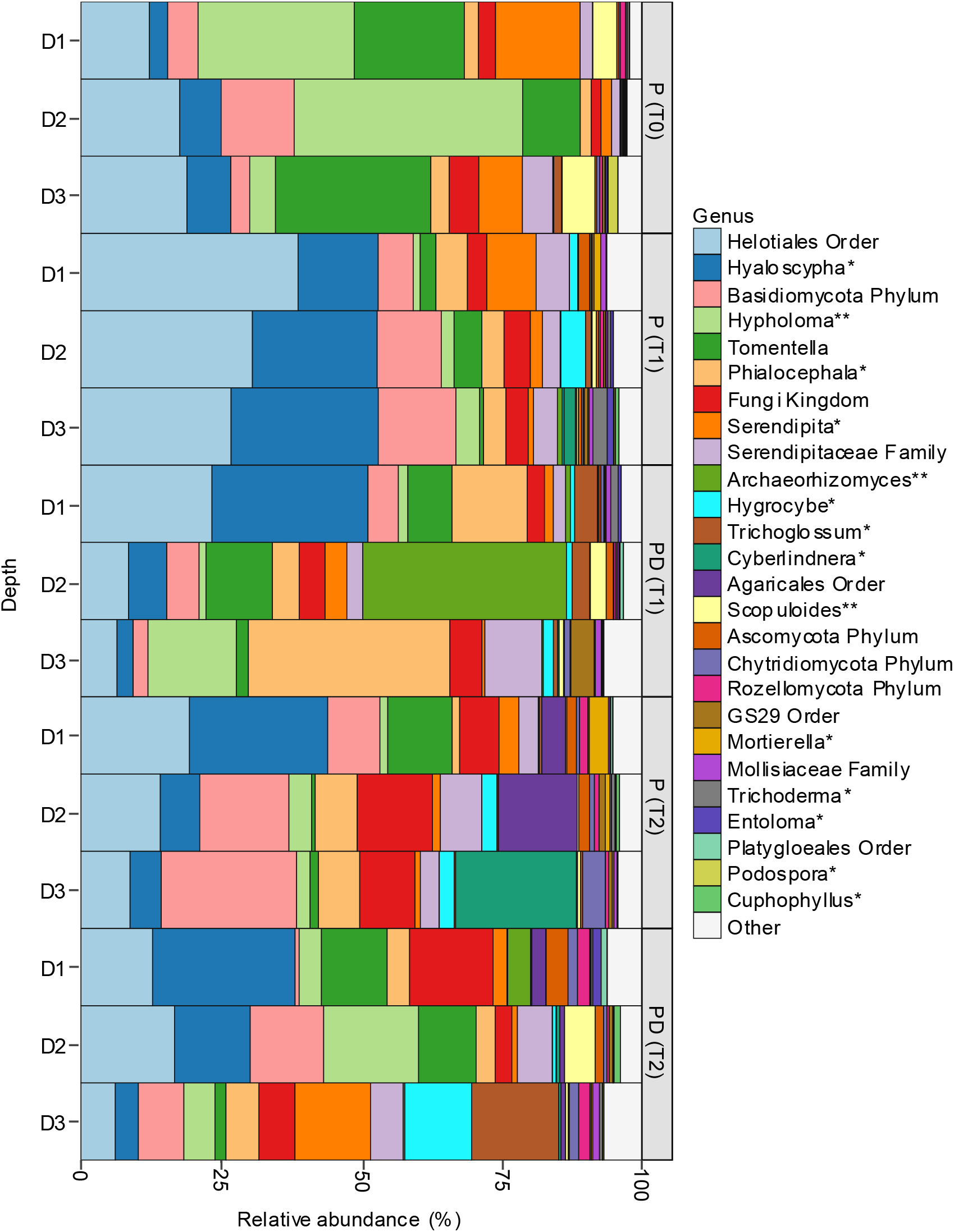
Relative abundance of fungal genera identified with ITS gene sequencing. Samples are organized by treatment (peat with dust [PD] and peat without dust [P]), timepoint (T0, T1, and T2 months), and peat depth (D1: 0-6, D2: 6-12, and D3: 12-18 cm). Fungal genera shown in the legend were prevalent in at least one sample and had a total relative abundance >1% when summed across all samples. Trophic mode classifications were retrieved from the FUNGUILD database (version 1.1). Genera labelled with one asterisk were classified under multiple trophic modes but included saprotrophy, while two asterisks indicate genera that were only classified with the saprotrophic lifestyle.

### 3.4. Peat pH

Changes to the pH of peat with the PD treatment were apparent in the upper zone of the peat. Peat pH_CaCl2_ was higher in D1 of the PD treatment when compared to the control after 2 months (P: pH = 3.72 ± 0.30; PD: pH = 4.66 ± 0.58) but showed little difference below the surface zone (Table 3). The pH_CaCl2_ in the control samples was generally stable between T0 and T2 months, indicating that mesocosm conditions did not notably influence peat pH.

**Table 3:** Peat pH_CaCl2_ measurements in the mineral dust treatment and control.

| Depth | Sample type | 0 months | 1 month | 2 months |
| --- | --- | --- | --- | --- |
| D1 | Control | $3.69 \pm 0.29$ | $3.60 \pm 0.04$ | $3.72 \pm 0.30$ |
| | Treatment | NA | $4.17 \pm 0.10$ | $4.66 \pm 0.58$ |
| D2 | Control | $4.00 \pm 0.22$ | $3.84 \pm 0.08$ | $3.97 \pm 0.05$ |
| | Treatment | NA | $4.14 \pm 0.12$ | $4.06 \pm 0.03$ |
| D3 | Control | $3.87 \pm 0.27$ | $4.14 \pm 0.16$ | $4.14 \pm 0.01$ |
| | Treatment | NA | $3.99 \pm 0.03$ | $4.12 \pm 0.01$ |

**Table 4:** Permutational analysis of variance (PERMANOVA) for fungal and bacterial communities with the Aitchison distance matrix (n permutations = 999). These results were consistent when count data was agglomerated to the genus or family taxonomic rank. Significant results (p < 0.05) are bolded and have an asterisk.

| Microbial group | Taxonomic Level | Parameter | Df | SumOfSqs | R<sup>2</sup> | F | p-value |
| --- | --- | --- | --- | --- | --- | --- | --- |
| Prokaryotes (16S rRNA gene) | Family | Treatment | 1 | 5508 | 0.04319 | 0.93 | 0.584 |
|  |  | **Time** | **2** | **15408** | **0.12083** | **1.30** | **0.047\*** |
|  |  | **Depth** | **2** | **15460** | **0.12124** | **1.30** | **0.034\*** |
|  |  | Treatment: Time | 1 | 5706 | 0.04475 | 1.00 | 0.401 |
|  |  | Treatment: Depth | 2 | 12457 | 0.09769 | 1.10 | 0.237 |
|  |  | Time: Depth | 4 | 23689 | 0.18577 | 1.04 | 0.328 |
| Fungi (ITS gene) | Genus | Treatment | 1 | 3726 | 0.07927 | 1.18 | 0.067 |
|  |  | **Time** | **2** | **7363** | **0.15667** | **1.16** | **0.028\*** |
|  |  | **Depth** | **2** | **7500** | **0.15958** | **1.19** | **0.015\*** |
|  |  | Treatment: Time | 1 | 3323 | 0.07070 | 1.09 | 0.208 |
|  |  | Treatment: Depth | 2 | 6708 | 0.14273 | 1.10 | 0.150 |
|  |  | Time: Depth | 4 | 12562 | 0.26728 | 1.03 | 0.384 |

### 3.5. Bioavailability of base cations and other metals

We assessed the general chemical properties of the prepared dust to identify particular elements that could influence microbial function. The dust had an average pH_CaCl2_ of 6.06 ± 0.45, with low amounts of total carbon (< 0.0015%) and nitrogen (<0.02%) (see Table S3). Iron occurred in the highest concentration out of all elements assessed (approximately 75000 mg/kg), followed by Al and Ca. Total phosphorus was low in the dust (approximately 660 mg/kg) compared to other tested elements (Table A.2), while orthophosphate was below detection.

Sequential chemical extraction provided insights into the bioavailability of elements in the peat after treatment with dust. We focused on base cations (Ca, Mg, and K), transition metals (Zn, Cu, Ni, Al, and Fe), and Al. Total concentrations of the elements were typically higher in the PD treatment compared to the control, but this depended on peat depth and soil fraction (Figure 5). The bioavailability of base cations and other metals was also generally higher in the PD treatment; this is supported by the increased concentrations of metals in the water soluble, exchangeable, and labile organics fractions. The relative distribution of tested elements across each soil fraction tended to be similar between the PD treatment and control after 2 months.

**Figure 5.**
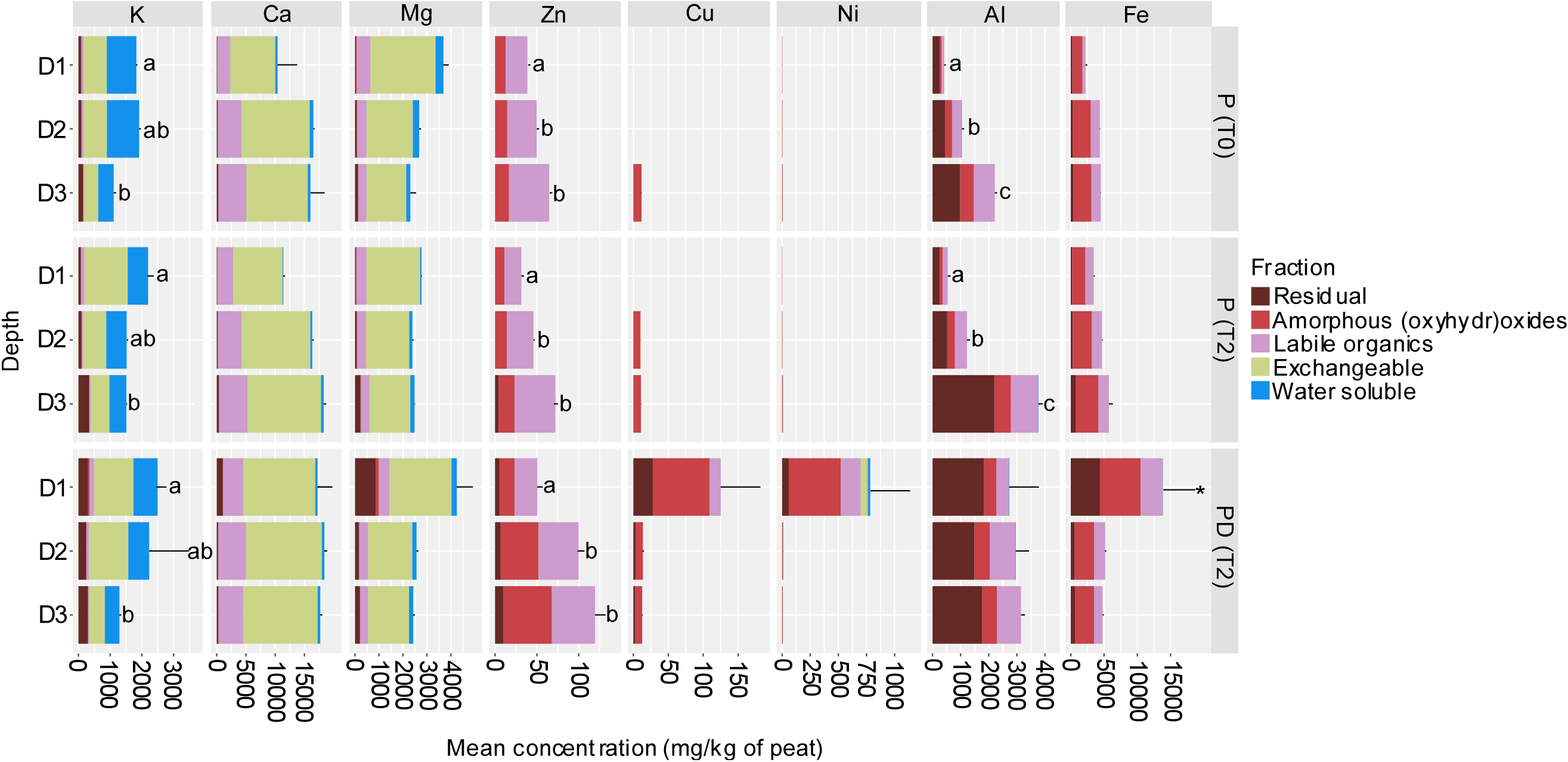
Mean concentrations of base cations and trace metals in peat samples (mg/kg of peat soil). Samples are organized by treatment (peat with dust [PD] and peat without dust [P]), timepoint (T0, and T2 months), and peat depth (D1: 0-6, D2: 6-12, and D3: 12-18 cm). The total element concentration is further organized by contributions from each tested soil fraction. Letters indicate significant differences in total element concentration (*i.e.,* summed across all fractions) between peat depths within the same treatment and timepoint (*i.e.,* a significant result from Kruskal-Wallis and Dunn’s post-hoc test). Asterisks indicate significant differences in total element concentration between the dust treatment and the respective control at the same timepoint and depth (Dunn’s test).

The PD treatment had higher total concentrations of base cation elements in D1 when compared to the control at T2 (Figure 5). The average concentration of exchangeable Ca across all depths was significantly higher in the PD treatment at T2 when compared to the control (Z_PD-P_ = 3.01, p < 0.01). The additional base cation content came mainly from the exchangeable fraction, which is consistent with the increased mobilization of these relatively soluble elements. The total concentrations of K, Ca, and Mg in the PD treatment were more similar to the control at depths D2 and D3 after two months.

Similar to the base cations, the total concentrations of Zn, Cu, Ni, Fe, and Al were higher in D1 of the treated peat soil compared to the control. The total Fe concentration was significantly higher in D1 relative to the control (Z_PD-P_ = 3.98, p < 0.05). Transition metals and Al were predominately located in the labile organics, amorphous (oxyhydr)oxide, and residual fractions (Figure 5). The vertical mobility and resulting fractionation at lower depths was variable between Zn, Cu, Ni, Fe, and Al. The enrichment of Fe from dust was largely restricted to D1, where the concentration of Fe was significantly higher than the control in both the labile organics and amorphous (oxyhydr)oxide fractions (p < 0.001). Aluminum, which was mainly associated with the residual fraction, was also higher in the treated peat in D1 and D2; this suggests that intact dust particles migrated down the column. Both Cu and Ni were higher in the most bioavailable fractions (water-soluble, exchangeable, and labile organics) after two months for the treated peat, but these elements did not notably migrate down the column. In the control samples, Cu and Ni were often below detection limits, except for a low concentration in the amorphous (oxyhydr)oxide fraction. Zinc was the most mobile metal, with dust application resulting in a higher content of Zn throughout the column compared to the control, mainly in the labile organics and amorphous (oxyhydr)oxide fractions (Figure 5).

Temporal trends in element concentrations for the control peat showed the natural variation amongst peat samples. The concentration of aluminum in the lowest zone of the control at T2 was around 70% higher relative to T0 samples (Figure 5). There was no evidence that Al migrated from the upper depths by T2, and most of the additional Al in D3 was located in the residual fraction. This result suggests natural variation in the mineral content of the peat rather than the leaching of Al down the column.

### 3.5. Phosphorus bioavailability

Phosphorus was of interest given that peatlands are typically low in this nutrient element. Most phosphorus was bound to amorphous (oxyhydr)oxides, which decreased over time in D1 and D2 of the PD treatment and the control (Figure 6). The total concentration of phosphorus was not significantly different between the PD treatment and control after 2 months. However, the total concentrations of P were 15% and 11% lower in D1 and D2 of the PD treatment relative to the control at T2.

**Figure 6.**
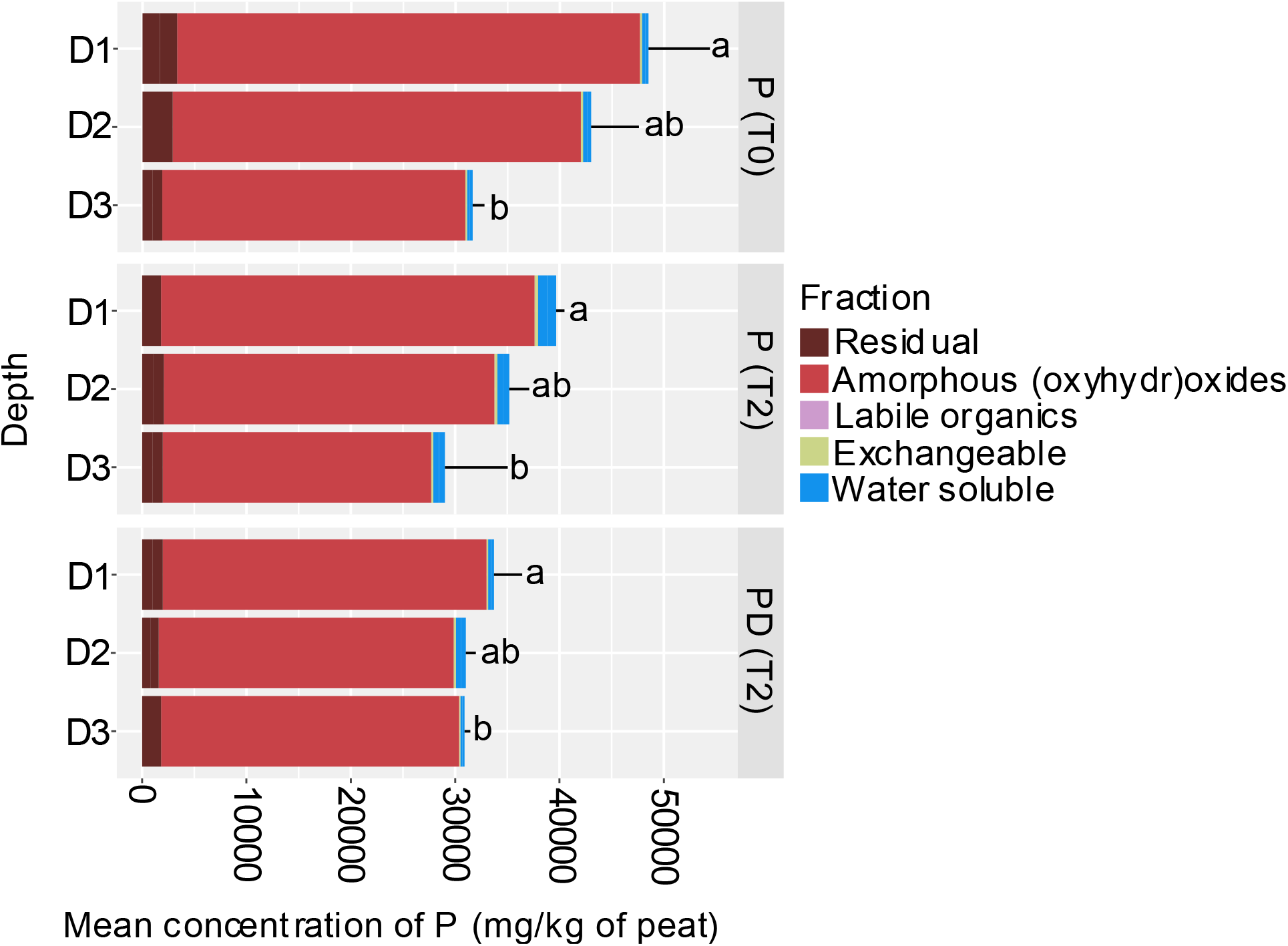
Mean concentrations (mg/kg of peat soil) of total phosphorus. Samples are organized by treatment (peat with dust [PD] and peat without dust [P]), timepoint (T0, and T2 months), and peat depth (D1: 0-6, D2: 6-12, and D3: 12-18 cm). The total element concentration is further organized by contributions from each tested soil fraction. Total concentrations with different letters indicate significant differences between peat depths within the same treatment and timepoint (*i.e.,* a significant result from Kruskal-Wallis and Dunn’s post-hoc test).

## 4. Discussion

### 4.1. Dust as a stimulant for microbial heterotrophy in peat

The application of dust induced diverse changes to the structure and activity of peatland microbial communities through time. The initial reduction in exoenzyme activity and capacity for mineralizing carbon substrates could explain the parallel increase in the relative abundances of fungal genera *Archaeorhizomyces* and *Phialocephala,* which are considered slow growing fungi that are resilient to high metal concentrations (Wang *et al.*, 2021; Haruma *et al.*, 2025).

The dust treatment may have induced a lag phase in the activity of the microbial community, which has been previously reported with changing nutrient or mineral regimes in soils (Zheng *et al.*, 2009; Wang *et al.*, 2016).

Several lines of evidence suggest that the dust treatment stimulated microbial heterotrophy. These include the higher carbon substrate use in the surface soil, increased potential activities of exoenzymes involved in C and N cycling within the middle depth, and changes to microbial community structure across the entire soil column. A feasible explanation for this response to the dust is that the release of minor macronutrients (e.g., Ca, Mg, K) and trace nutrients (e.g., Fe, Zn, Cu, Ni) stimulated the production of cellular structures and redox reactions involving C, N, and S (Karaffa *et al.*, 2021; Dai *et al.*, 2023). For example, Cu is incorporated into oxidases found among microbial communities in ombrotrophic peat and has been reported to stimulate CH_4_ oxidation and N_2_ fixation in methanotrophs from peat moss (Lin et al., 2014). Other metals including Fe, Zn, and Ca are also involved in later stages of aerobic methanotropy (Glass and Orphan, 2012). Previous experiments have demonstrated that cation amendments, such as Cu^2+^ and Ca^2+^, could increase CH_4_ respiration by methanogens in deep peat (Basiliko and Yavitt, 2001; Thomas and Pearce, 2004). In contrast, Keller and Wade (2018) reported a slight, but consistent inhibition of anaerobic respiration in peat with Cu, Fe, or Ni amendments. The short incubation time used in that study (one month) and the application of metals in dissolved form could have led to a lag phase, as we also observed. Besides directly stimulating microbial processes, an influx of dissolved base cations from the dust could have released pre-existing exoenzymes sorbed to soil surfaces through exchange processes, as previously observed for *Sphagnum* dominated peat (Gogo and Pearce, 2009; Gogo *et al.*, 2010). The higher relative abundance of putative saprotrophic fungi in the dust-treated peat also aligns with the observed stimulation in microbial activity. Saprotrophs are characterized by a high dependence on the enzyme-mediated breakdown of organic matter for energy and nutrients (Thormann, 2006). Little is known about the cellular requirements of metallic nutrient elements that support different fungal guilds. Previous observations by Peltoniemi *et al.* (2021) demonstrated that soil iron content was a strong predictor of fungal community structure in boreal peatland forests. Metal nutrient elements may support the production of exoenzymes that are frequently released by saprotrophs, such as Cu-dependent laccases (Fonseca *et al.*, 2010; Janusz *et al.*, 2017), or the stabilization of amylases by Ca (Nazmi *et al.*, 2006). However, the interactions between metal cofactors and exoenzymes remain poorly understood in comparison to intracellular enzymes (He *et al.*, 2023).

Acid phosphatase was the only exoenzyme we observed that decreased in activity throughout the soil exposed to dust. This could indicate that limitations of the major macronutrient phosphorus were relieved with the dust treatment, which could have stimulated the microbial mineralization of organic matter (Cui *et al.*, 2023). The trapping of phosphorus by association with metal oxides is considered an important limiting factor for bog productivity (Shang *et al.*, 1996; Zhao *et al.*, 2021). However, because the peat samples were incubated on Ecoplates^TM^ in phosphate buffered saline solution, a reduction in phosphorus limitation in the peat does not fully explain the enhanced carbon substrate mineralization observed in the surface zone.

Our results suggest that the solid and dissolved components of the dust reduced the bioavailability of dissolved organic matter (DOM) across the soil profile. A substantial fraction of the increased transition metals and Al provided by the dust were associated with the labile organics fraction by two months. With very low organic matter content in the mineral dust, a majority of the metal enrichments in the labile organics phase likely arose from the complexation of peat organic matter with metals released from mineral oxides within the dust. Both humic and fulvic acids in *Sphagnum*-dominated peat are efficient scavengers of metals bound in iron (oxyhydr)oxides (Zhao *et al.*, 2021; Butt *et al.*, 2024). Mineral oxide surfaces and metal ions are well known to complex and aid in the polymerization of OM (Zhao *et al.*, 2019). A reduction in the bioavailability of DOM could incentivize microbes to allocate more energy towards degrading polymerized substrates, which would explain the observed increase in the carbohydrate polymers glycogen and α-cyclodextrin. Additionally, the stimulation of β-xylosidase activity in the middle depth supports that microbial communities were in the early stages of recalcitrant carbohydrate biodegradation (Novak *et al.*, 2024). Because polymeric degradation is energetically and nutritionally expensive for microbes, higher proportions of carbon could be allocated towards exoenzyme production and cellular respiration compared to more labile carbon sources (Dang *et al.*, 2024). Conversely, organic carbon may be protected against degradation through the association with dust minerals.

### 4.2. Spatial variation in metal enrichments and microbial responses to dust inputs

Our results show that chemical and biological parameters throughout the peat column changed when dust was applied. The nature of these impacts and implications for carbon mineralization was dependent on the peat depth. Despite having the highest average growth rate during the C mineralization experiments, microbial communities in the surface zone of the treated peat did not experience a parallel increase in the activity of C or N cycling exoenzymes. The relative abundance of putative oligotrophic bacterial families was also higher in the surface zone (e.g., Acidobacteriaceae (Subgroup 1) and Koribacteraceae), which suggests that activity was not stimulated *in situ* despite the increases in some nutrient elements (Elliott *et al.*, 2015; Morrison *et al.*, 2021). Exoenzyme activity could have also been suppressed at the surface due to the increase in the bioavailability of elements that are toxic at low concentrations. Of the transition metals we assessed, Ni was the most bioavailable with the dust treatment. Free Ni ions can inhibit both intracellular and extracellular enzymes by acting as allosteric inhibitors, binding to active sites, and replacing metals in metalloenzymes (Lankinen *et al.*, 2011). Another pathway for the reduction in exoenzyme activity at the surface depth is the increased sorption and subsequent inhibition of exoenzymes by the mineral material added to the surface (Vepsäläinen, 2001; Sheng *et al.* 2023). Over longer time periods, these sorbed exoenzymes may be protected from proteolytic degradation, be released over time, and help to hydrolyze organic matter (Olagoke *et al.*, 2020). Carbon substrate utilization was nevertheless highest in the surface zone of treated peat. A plausible explanation for the discrepancy is that peat microbial communities are removed from the soil matrix and grown in a buffer solution for the Ecoplate^TM^ incubations.

Based on the smaller enrichment of elements bound to residual fractions in the middle zone, we suspect that much of the dust remained in the surface zone, given its particulate nature and the low permeability of peat. A lower amount of particulates from the dust entering the middle zone could have lowered the degree of exoenzyme sorption to mineral surfaces compared to the upper zone. To investigate this further, future work should characterize mineral content with depth after exposure to dust deposition. In the deepest zone, we observed no clear increases in the activity of soil microbes with the dust treatment. This pattern aligns with the relatively small chemical changes observed in this zone. The lowest zone was also permanently saturated, where lower oxygen concentrations are likely an important variable limiting functional responses from microbial communities to the dust treatment.

### 4.3. Site specific considerations for interpreting peatland microbial responses to dust inputs

In our study, we used a rather potent exposure of dust, comparable to one year of deposition based on estimates in bogs near the Athabasca oil sands in Alberta, Canada (Mullan-Boudreau *et al.*, 2017) and the Great Hinggan Mountain region in northeast China (Bao *et al.*, 2012). The potential for mineral dusts to release essential elements into peat environments will vary as a function of deposition rate, mineral composition and particle size. However, the few available estimates of dust loading to northern peatlands do not consider the weathering of dust minerals after deposition. Our assessment of changes in microbial activity after introducing dust to peat soil may be particularly relevant for peatlands near large extraction or other land development, or near dry regions. Once deposited, the type of minerals, their solubilities, element speciation in soil pore water, interactions with solid phases, and metal redox reactions will influence the bioavailability of metals released during dust weathering (Le Roux *et al.*, 2006; Linke and Gislason, 2018). The dust we applied contained metal oxides that could release metals like Fe, Ni and Zn through acidic and reductive dissolution (Olsson and Wallander, 1998; Grand and Lavkulich, 2015). The major mineral constituents, silicate (i.e., albite and actinolite) and aluminosilicate minerals (i.e. clinochlore and biotite) (Hettinga *et al.*, in press) will release some nutrient elements during weathering but contain relatively low concentrations compared to the aluminosilicates and metal oxides, and weather more slowly.

Microbial responses to elements released into bioavailable fractions will likely vary based on the unique chemical context of the peatland. The natural peat soil used in this study was most representative of an ombrotrophic bog, with low total concentrations of Fe, Al, Cu, Ni, and Zn as observed in bogs globally (Osborne *et al.*, 2024). However, concentrations of Ca and Mg in the most bioavailable element fractions were similar to those in moderate fens (Lipatov *et al.*, 2018); which can be attributed to the calcite and dolomite-rich geology of the region (Coniglio *et al.*, 2003). Changes in the nutrient budgets of ombrotrophic bogs have also been shown to not only increase organic matter breakdown, but also to alter plant species composition (Bubier *et al.*, 2007). Therefore, assessments of organic matter bioavailability, microbial mineralization, and above-ground productivity over a longer period and in the field are the next steps for understanding how dust deposition could influence carbon stability in peatlands.

## 5. Conclusions

In this study, we investigated the extent to which dust inputs could weather in ombrotrophic peat and influence the functioning of subsurface microbial communities. Our study represents a first look at how minerals which could conceivably exist in atmospheric dust can alter microbial activity in peat soil through the release of multiple metallic elements. By using an extended sequential extraction approach, we show that mineral dust can readily release metals into the most bioavailable soil fractions including water-soluble, exchangeable, and labile organics. Although the enrichment of metals after dust application decreased with depth, zinc was particularly mobile throughout the column when complexed with labile organic molecules. Responses from microbial communities to dust amendment were highly dependent on the unique chemical perturbations that occurred at each depth. Microbial communities closest to the site of dust application showed the highest capacity to mineralize carbon substrates on Biolog Ecoplates^TM^. However, the lack of stimulation in exoenzyme activity indicates that dust may reduce microbial activity near dust particles. In the middle depth, the activities of β-glucosidase, β-xylosidase, and NAGase were stimulated by the dust treatment. Throughout the core we observed increases in the representation of several saprotrophic fungal genera (e.g., *Tomentella*, *Hypholoma*, *Serendipita*, and more) that are characterized by high nutrient demands and are specialized for the enzyme-mediated breakdown of organic matter. A higher capacity to degrade carbohydrate polymers throughout the core suggests that the mineral and metal content from dust had reduced the availability of dissolved organic matter and incentivized microbial degradation of recalcitrant carbon sources. This work supports that dust deposition has the potential to moderate the microbial mineralization of organic carbon in ombrotrophic peatlands.

## CRediT authorship contribution statement

**Jordan Thakar**: Conceptualization, Methodology, Investigation, Data Curation, Visualization, Formal Analysis, Writing-Original Draft, Writing-Review & Editing. **E. Kathryn Hettinga**: Conceptualization, Methodology, Formal Analysis, Investigation, Data Curation, Writing-Review & Editing, Visualization. **Kimber E. Munford**: Conceptualization, Methodology, Formal Analysis, Investigation, Data Curation, Writing-Review & Editing, Visualization, Supervision. **Susan Glasauer**: Conceptualization, Methodology, Formal Analysis, Investigation, Resources, Project Administration, Data Curation, Writing-Review & Editing, Visualization, Supervision.

## Data availability

Chemical and biological data from this study are deposited in the Open Science Framework available at: osf.io/6jyb2. Raw sequencing data from this study is deposited in NCBI’s Sequencing Read Archive (Submission ID: SUB16346348; BioProject ID: PRJNA1498746). R scripts for processing and analyzing DNA sequencing data are available on GitHub: https://github.com/jthaka/Cranberry_Bog_Dust_Experiment.git

## Acknowledgements

We would like to also thank Peter Smith at the University of Guelph for conducting the ICP-OES analysis. Also, we would like to thank the Physics Department Workshop at the University of Guelph for assisting in the design and construction of our mesocosm apparatus.

