## Supplemental Tables for "Mineral dust stimulates microbial exoenzyme activity and enhances carbon mineralization capabilities in nutrient-poor peat soil"

### Table S1- Baseline soil and porewater chemistry in Cranberry Bog. Peat samples were retrieved on October 27, 2022. Porewater measurements were performed in-field.

| Depth | TC (%) | IC (%) | OC (%) | TN (%) | pH_CaCl2_ | Temperature (℃) | Conductivity (µS/cm) | Dissolved oxygen (%) |
| --- | --- | --- | --- | --- | --- | --- | --- | --- |
| 0-6 cm | 45.4 | 0.107 | 45.3 | 1.52 | 3.69 ± 0.24 | NA | NA | NA |
| 6-12 cm | 47.8 | 0.133 | 47.6 | 1.23 | 4.00 ± 0.18 | NA | NA | NA |
| 12-18 cm | 48.6 | 0.133 | 48.5 | 1.95 | 3.87 ± 0.22 | NA | NA | NA |
| Porewater | NA | NA | NA | NA | 4.2 ± 0.02 | 9.5 ± 0.15 | 112.4 µS/cm ± 4.5 | 6.3% ± 0.9 |

### Table S2- Total element concentrations (mg/kg) in mine tailings (<20 µm fraction size) extracted by the reverse aqua regia protocol and measured with ICP-OES. All values represent the average concentration from eight extraction replicates.

|  | Fe | Al | Ca | Mg | S | K | Ni | P | Mn | Cu | Zn |
| --- | --- | --- | --- | --- | --- | --- | --- | --- | --- | --- | --- |
| >20 µm tailings dust | 75006.26 ± 2834.10 | 21736.87 ± 626.99 | 18981.09 ± 446.50 | 12273.083 ± 39.92 | 12400.06 ± 840.24 | 3736.20 ± 94.08 | 1945.60 ± 53.88 | 659.31 ± 13.38 | 569.64 ± 8.01 | 481.87 ± 33.23 | 100.79 ± 1.38 |

### Table S3- General chemistry of original mine tailings material that was used for mesocosm experiments (prior to sieving for the <20 µm fraction).

|  | pH | Total carbon (%) | Inorganic carbon (%) | Organic carbon (%) | Total nitrogen (%) |
| --- | --- | --- | --- | --- | --- |
| Tailings material | 6.06 ± 0.45 | 0.00151 ± 2.20 | 0.000443 ± 1.28 | 0.00106 ± 3.56 | <0.02 |

### Table S4- Total element enrichment (%) between the treated peat and control. Enrichment is based on the % difference between the mean concentrations (n = 5 replicates) between the treatment and control. Null values for Cu and Ni are a result of the mean concentrations in the control being below detection.

|  | Soil depth | | |
| --- | --- | --- | --- |
| Element | D1 | D2 | D3 |
| K | 14.63 | 46.09 | -14.70 |
| Ca | 51.73 | 12.72 | -3.23 |
| Mg | 53.46 | 7.18 | -2.33 |
| Zn | 57.06 | 114.64 | 69.70 |
| Cu | NA | 27.57 | 16.50 |
| Ni | NA | 142.24 | -9.42 |
| Al | 403.06 | 141.33 | -16.39 |
| Fe | 302.44 | 9.34 | -16.65 |

### Table S5- Results from Shapiro-Wilks tests at p < 0.05 for each chemical and biological dataset which include metals and phosphorus concentrations, activity of exoenzymes, and absorbance on Biolog Ecoplates^TM^.

| Data | Subset | n samples | W statistic | P value |
| --- | --- | --- | --- | --- |
| Total element concentration | K | 44 | 0.812 | 5.46E-06 |
|  | Ca | 44 | 0.787 | 1.57E-06 |
|  | Mg | 44 | 0.761 | 4.55E-07 |
|  | Zn | 44 | 0.880 | 2.79E-4 |
|  | Cu | 47 | 0.464 | 1.82E-11 |
|  | Ni | 44 | 0.340 | 8.33E-13 |
|  | Al | 44 | 0.898 | 9.39E-4 |
|  | Fe | 44 | 0.656 | 6.70E-09 |
|  | P | 44 | 0.915 | 3.17E-3 |
|  | PO_4_^3-^ | 45 | 0.615 | 1.25E-09 |
| Exoenzyme activity | Acid phosphatase | 75 | 0.947 | 3.63E-03 |
|  | β-Glucosidase | 75 | 0.979 | 2.55E-01 |
|  | β-Xylosidase | 75 | 0.869 | 1.43E-06 |
|  | NAGase | 75 | 0.688 | 2.41E-11 |
| Absorbance on Biolog Ecoplates^TM^ | Total | 45 | 0.886 | 3.61E-04 |
|  | Amines | 45 | 0.856 | 5.33E-05 |
|  | Amino Acids | 45 | 0.901 | 1.03E-03 |
|  | Carbohydrates | 45 | 0.882 | 2.64E-04 |
|  | Carboxylic acids | 45 | 0.878 | 2.14E-04 |
|  | Phenolics | 45 | 0.810 | 3.93E-06 |
|  | Polymers | 45 | 0.895 | 6.87E-04 |

### Table S6- Significant (p < 0.05) Kruskal-Wallis and post-hoc Dunn’s test results for pairwise comparisons of mean element concentration between treatment and timepoint groups. Groups used for comparisons include the control at T0 (P_T0), no dust control at T2 months (P_T2), and the dust treatment after T2 months (PD_T2). Test results are organized by the corresponding element and soil fraction included in the assessment. Significant p-values were corrected with the Bonferroni correction.

|  |  |  |  | Dunn’s post hoc result | |
| --- | --- | --- | --- | --- | --- |
| Soil fraction | Element | KW | Comparison | Z statistic | p-value^BF^ |
| Water | Al | χ^2^(2) = 6.4, p = 0.0409, ε^2^ = 0.291 | P_T2 - PD_T2 | -2.51 | 0.0359 |
|  | Ca | χ^2^(2) = 8.67, p = 0.0131, ε^2^ = 0.202 | P_T0 - P_T2 | 2.94 | 0.0097 |
|  | K | χ^2^(2) = 7.6, p = 0.0224, ε^2^ = 0.177 | P_T0 - P_T2 | 2.59 | 0.0291 |
|  | Mg | χ^2^(2) = 18.96, p = 1e-04, ε^2^ = 0.441 | P_T0 - P_T2 | 4.32 | 0.0000 |
|  |  |  | P_T2 - PD_T2 | -2.59 | 0.0291 |
|  | P | χ^2^(2) = 29.19, p = 0, ε^2^ = 0.679 | P_T0 - P_T2 | -4.38 | 0.0000 |
|  |  |  | P_T2 - PD_T2 | 4.90 | 0.0000 |
| Exchangeable | Ca | χ^2^(2) = 22.18, p = 0, ε^2^ = 0.528 | P_T0 - PD_T2 | -4.62 | 0.0000 |
|  |  |  | P_T2 - PD_T2 | -3.01 | 0.0078 |
|  | Fe | χ^2^(2) = 6.23, p = 0.0443, ε^2^ = 0.249 | P_T2 - PD_T2 | -2.41 | 0.0485 |
|  | P | χ^2^(2) = 19.05, p = 1e-04, ε^2^ = 0.443 | P_T0 - P_T2 | -2.57 | 0.0306 |
|  |  |  | P_T2 - PD_T2 | 4.34 | 0.0000 |
| Labile organics | Al | χ^2^(2) = 11.94, p = 0.0026, ε^2^ = 0.278 | P_T0 - PD_T2 | -3.44 | 0.0017 |
|  | Fe | χ^2^(2) = 12.51, p = 0.0019, ε^2^ = 0.291 | P_T0 - PD_T2 | -3.52 | 0.0013 |
| amorphous (oxyhydr)oxides | Al | χ^2^(2) = 12.82, p = 0.0016, ­ε^2^ = 0.298 | P_T0 - PD_T2 | -3.58 | 0.0010 |
|  | Fe | χ^2^(2) = 18.04, p = 1e-04, ε^2^ = 0.419 | P_T0 - PD_T2 | -4.25 | 0.0001 |
|  | Mg | χ^2^(2) = 7.55, p = 0.0229, ε^2^ = 0.176 | P_T0 - P_T2 | 2.73 | 0.0192 |
|  | Zn | χ^2^(2) = 22.52, p = 0, ε^2^ = 0.524 | P_T0 - PD_T2 | -4.16 | 0.0001 |
|  |  |  | P_T2 - PD_T2 | -4.04 | 0.0002 |
|  | P | χ^2^(2) = 14.42, p = 7e-04, ε^2^ = 0.335 | P_T0 - PD_T2 | 3.80 | 0.0004 |
| Residual | Al | χ^2^(2) = 14.49, p = 7e-04, ε^2^ = 0.337 | P_T0 - PD_T2 | -3.79 | 0.0005 |
|  | Ca | χ^2^(2) = 11.68, p = 0.0029, ε^2^ = 0.278 | P_T0 - PD_T2 | -3.35 | 0.0025 |
|  | Fe | χ^2^(2) = 17.41, p = 2e-04, ε^2^ = 0.405 | P_T0 - PD_T2 | -3.91 | 0.0003 |
|  |  |  | P_T2 - PD_T2 | -3.18 | 0.0044 |
|  | K | χ^2^(2) = 14.11, p = 9e-04, ε^2^ = 0.344 | P_T0 - PD_T2 | -3.74 | 0.0006 |
|  | Mg | χ^2^(2) = 17.4, p = 2e-04, ε^2^ = 0.414 | P_T0 - PD_T2 | -4.07 | 0.0001 |
|  |  |  | P_T2 - PD_T2 | -2.78 | 0.0163 |
|  | P | χ^2^(2) = 15.91, p = 4e-04, ε^2^ = 0.37 | P_T0 - PD_T2 | 3.99 | 0.0002 |
| Total | Al | χ^2^(2) = 12.1, p = 0.0024, ε^2^ = 0.281 | P_T0 - PD_T2 | -3.47 | 0.0015 |
|  | Ca | χ^2^(2) = 13.01, p = 0.0015, ε^2^ = 0.303 | P_T0 - PD_T2 | -3.56 | 0.0011 |
|  | Fe | χ^2^(2) = 18.62, p = 1e-04, ­ε^2^ = 0.433 | P_T0 - P_T2 | -2.50 | 0.0371 |
|  |  |  | P_T0 - PD_T2 | -4.30 | 0.0001 |
|  | Zn | χ^2^(2) = 12.19, p = 0.0022, ε^2^ = 0.284 | P_T0 - PD_T2 | -2.75 | 0.0180 |
|  |  |  | P_T2 - PD_T2 | -3.23 | 0.0038 |
|  | P | χ^2^(2) = 13.82, p = 0.001, ε^2^ = 0.321 | P_T0 - PD_T2 | 3.72 | 0.0006 |

### Table S7- Significant (p < 0.05) Kruskal-Wallis and post-hoc Dunn’s test results for pairwise comparisons of mean element concentration between unique treatment-depth groupings. Treatments included the control at T2 months (P), and the dust treatment at T2 months (PD). Depths included the upper (D1), middle (D2), and lower (D3) peat layers. Test results are organized by the corresponding element and soil fraction included in the assessment.

|  |  |  |  | Corrected p-value (Bonferroni) | |
| --- | --- | --- | --- | --- | --- |
| Soil fraction | Element | KW | Comparison | Z statistic | p-value^BF^ |
| Water soluble | Ca | χ^2^(8) = 40.38, p = 0, eps2 = 0.939 | P_D2 - P_D1 | 4.80 | 0.0001 |
|  |  |  | P_D2 - P_D2 | 4.16 | 0.0011 |
|  |  |  | P_D1 - P_D3 | -4.06 | 0.0018 |
|  |  |  | P_D2 - P_D3 | -3.42 | 0.0224 |
|  |  |  | P_D2 - PD_D1 | 3.47 | 0.0187 |
|  | K | χ^2^(8) = 41.13, p = 0, eps2 = 0.957 | P_D1 - P_D3 | 3.48 | 0.0179 |
|  |  |  | P_D2 - P_D3 | 4.06 | 0.0018 |
|  |  |  | P_D2 - P_D3 | 3.69 | 0.0080 |
|  |  |  | P_D1 - PD_D3 | 4.14 | 0.0013 |
|  |  |  | P_D2 - PD_D3 | 4.75 | 0.0001 |
|  |  |  | PD_D1 - PD_D3 | 3.52 | 0.0155 |
|  | Mg | χ^2^(8) = 40.53, p = 0, eps2 = 0.943 | P_D1 - P_D1 | 4.80 | 0.0001 |
|  |  |  | P_D2 - P_D1 | 4.19 | 0.0010 |
|  |  |  | P_D1 - P_D2 | 4.19 | 0.0010 |
|  |  |  | P_D2 - P_D2 | 3.57 | 0.0129 |
|  |  |  | P_D1 - PD_D1 | -3.57 | 0.0129 |
|  | P | χ^2^(8) = 42.12, p = 0, eps2 = 0.98 | P_D3 - P_D1 | -4.00 | 0.0022 |
|  |  |  | P_D1 - PD_D1 | 3.69 | 0.0080 |
|  |  |  | P_D1 - PD_D3 | 4.80 | 0.0001 |
|  |  |  | P_D2 - PD_D3 | 3.94 | 0.0029 |
|  |  |  | P_D3 - PD_D3 | 3.82 | 0.0049 |
| Exchangeable | Ca | χ^2^(8) = 36.7, p = 0, eps2 = 0.874 | P_D1 - P_D3 | -3.48 | 0.0182 |
|  |  |  | P_D1 - PD_D2 | -4.21 | 0.0009 |
|  |  |  | P_D1 - PD_D2 | -3.90 | 0.0034 |
|  |  |  | P_D1 - PD_D3 | -3.67 | 0.0088 |
|  |  |  | P_D1 - PD_D3 | -3.32 | 0.0319 |
|  | Fe | χ^2^(8) = 14.03, p = 0.0293, eps2 = 0.561 | P_D3 - PD_D1 | -3.21 | 0.0283 |
|  | K | χ^2^(8) = 36.87, p = 0, eps2 = 0.857 | P_D3 - P_D1 | -4.21 | 0.0009 |
|  |  |  | P_D1 - P_D3 | 3.45 | 0.0204 |
|  |  |  | P_D3 - PD_D1 | -3.86 | 0.0041 |
|  |  |  | P_D1 - PD_D3 | 4.19 | 0.0010 |
|  |  |  | PD_D1 - PD_D3 | 3.82 | 0.0049 |
|  | Mg | χ^2^(8) = 38.44, p = 0, eps2 = 0.894 | P_D1 - P_D3 | 3.78 | 0.0057 |
|  |  |  | P_D1 - P_D3 | 4.04 | 0.0019 |
|  |  |  | P_D3 - PD_D1 | -3.48 | 0.0183 |
|  |  |  | P_D3 - PD_D1 | -3.72 | 0.0072 |
|  |  |  | P_D1 - PD_D3 | 3.87 | 0.0040 |
|  |  |  | PD_D1 - PD_D3 | 3.55 | 0.0141 |
|  | P | χ^2^(8) = 37.01, p = 0, eps2 = 0.861 | P_D1 - PD_D1 | 3.99 | 0.0024 |
|  |  |  | P_D2 - PD_D1 | 3.37 | 0.0268 |
|  |  |  | P_D1 - PD_D3 | 4.80 | 0.0001 |
|  |  |  | P_D2 - PD_D3 | 4.19 | 0.0010 |
| Labile organics | Al | χ^2^(8) = 42.06, p = 0, eps2 = 0.978 | P_D1 - P_D3 | -4.80 | 0.0001 |
|  |  |  | P_D2 - P_D3 | -3.57 | 0.0129 |
|  |  |  | P_D1 - P_D3 | -4.19 | 0.0010 |
|  |  |  | P_D1 - PD_D2 | -4.16 | 0.0011 |
|  |  |  | P_D1 - PD_D2 | -3.55 | 0.0141 |
|  |  |  | P_D1 - PD_D3 | -3.59 | 0.0117 |
|  | Ca | χ^2^(8) = 40.44, p = 0, eps2 = 0.941 | P_D1 - P_D3 | -3.89 | 0.0036 |
|  |  |  | P_D3 - P_D1 | 3.31 | 0.0339 |
|  |  |  | P_D1 - P_D3 | -4.33 | 0.0005 |
|  |  |  | P_D1 - P_D3 | -3.72 | 0.0072 |
|  |  |  | P_D1 - PD_D2 | -4.26 | 0.0007 |
|  |  |  | P_D1 - PD_D2 | -3.64 | 0.0097 |
|  | Fe | χ^2^(8) = 41.32, p = 0, eps2 = 0.961 | P_D1 - P_D3 | -3.87 | 0.0040 |
|  |  |  | P_D1 - PD_D1 | -4.80 | 0.0001 |
|  |  |  | P_D1 - PD_D3 | -4.06 | 0.0018 |
|  |  |  | P_D1 - PD_D2 | -3.87 | 0.0040 |
|  |  |  | PD_D1 - PD_D3 | 3.47 | 0.0187 |
|  | K | χ^2^(8) = 37.48, p = 0, eps2 = 0.872 | P_D3 - P_D1 | -3.69 | 0.0081 |
|  |  |  | P_D3 - PD_D1 | -4.29 | 0.0006 |
|  |  |  | P_D3 - PD_D1 | -3.62 | 0.0107 |
|  |  |  | P_D1 - PD_D3 | 3.62 | 0.0107 |
|  |  |  | PD_D1 - PD_D3 | 4.26 | 0.0007 |
|  | Mg | χ^2^(8) = 38.89, p = 0, eps2 = 0.904 | P_D1 - P_D3 | 3.66 | 0.0092 |
|  |  |  | P_D1 - P_D3 | 3.52 | 0.0155 |
|  |  |  | P_D1 - PD_D3 | 4.73 | 0.0001 |
|  |  |  | P_D1 - PD_D3 | 3.50 | 0.0170 |
|  |  |  | PD_D1 - PD_D3 | 4.11 | 0.0014 |
|  | Zn | χ^2^(8) = 39.63, p = 0, eps2 = 0.922 | P_D3 - P_D1 | 3.54 | 0.0144 |
|  |  |  | P_D1 - P_D3 | -3.89 | 0.0036 |
|  |  |  | P_D1 - PD_D2 | -3.61 | 0.0112 |
|  |  |  | P_D1 - PD_D3 | -3.58 | 0.0123 |
|  |  |  | P_D1 - PD_D3 | -4.47 | 0.0003 |
|  |  |  | PD_D1 - PD_D3 | -3.48 | 0.0178 |
| Amorphous (oxyhydr)oxides | Al | χ^2^(8) = 42.2, p = 0, eps2 = 0.981 | P_D1 - P_D3 | -4.78 | 0.0001 |
|  |  |  | P_D2 - P_D3 | -3.52 | 0.0155 |
|  |  |  | P_D1 - P_D3 | -4.16 | 0.0011 |
|  |  |  | P_D1 - PD_D2 | -4.14 | 0.0013 |
|  |  |  | P_D1 - PD_D2 | -3.52 | 0.0155 |
|  |  |  | P_D1 - PD_D3 | -3.64 | 0.0097 |
|  | Ca | χ^2^(8) = 34.66, p = 0, eps2 = 0.806 | P_D1 - P_D3 | 3.46 | 0.0193 |
|  |  |  | P_D1 - P_D2 | 3.27 | 0.0381 |
|  |  |  | P_D1 - P_D3 | 4.23 | 0.0008 |
|  |  |  | P_D1 - P_D3 | 3.82 | 0.0049 |
|  | Fe | χ^2^(8) = 36.78, p = 0, eps2 = 0.855 | P_D1 - P_D3 | -3.59 | 0.0117 |
|  |  |  | P_D1 - PD_D1 | -4.80 | 0.0001 |
|  |  |  | P_D1 - PD_D3 | -4.19 | 0.0010 |
|  |  |  | P_D1 - PD_D2 | -3.35 | 0.0293 |
|  | K | χ^2^(8) = 25.57, p = 0.0012, eps2 = 0.673 | P_D2 - PD_D1 | -3.40 | 0.0245 |
|  |  |  | P_D3 - PD_D1 | -4.28 | 0.0007 |
|  |  |  | PD_D1 - PD_D3 | 3.35 | 0.0290 |
|  | Mg | χ^2^(8) = 37.57, p = 0, eps2 = 0.874 | P_D1 - P_D1 | 3.80 | 0.0051 |
|  |  |  | P_D1 - PD_D1 | -4.42 | 0.0004 |
|  |  |  | P_D1 - PD_D3 | 3.93 | 0.0031 |
|  |  |  | PD_D1 - PD_D3 | 4.54 | 0.0002 |
|  | Zn | χ^2^(8) = 38.87, p = 0, eps2 = 0.904 | P_D1 - PD_D2 | -3.45 | 0.0204 |
|  |  |  | P_D1 - PD_D2 | -3.96 | 0.0027 |
|  |  |  | P_D1 - PD_D3 | -4.01 | 0.0022 |
|  |  |  | P_D2 - PD_D3 | -3.25 | 0.0416 |
|  |  |  | P_D1 - PD_D3 | -4.53 | 0.0002 |
|  |  |  | P_D2 - PD_D3 | -3.47 | 0.0187 |
|  | P | χ^2^(8) = 35.62, p = 0, eps2 = 0.828 | P_D1 - P_D3 | 3.79 | 0.0054 |
|  |  |  | P_D2 - P_D3 | 3.25 | 0.0416 |
|  |  |  | P_D1 - PD_D2 | 3.89 | 0.0036 |
|  |  |  | P_D2 - PD_D2 | 3.35 | 0.0293 |
|  |  |  | P_D1 - PD_D3 | 3.72 | 0.0072 |
| Residual | Al | χ^2^(8) = 40.49, p = 0, eps2 = 0.942 | P_D1 - P_D3 | -4.53 | 0.0002 |
|  |  |  | P_D2 - P_D3 | -3.42 | 0.0224 |
|  |  |  | P_D1 - P_D3 | -4.46 | 0.0003 |
|  |  |  | P_D1 - PD_D1 | -3.57 | 0.0129 |
|  |  |  | P_D1 - PD_D1 | -3.50 | 0.0170 |
|  |  |  | P_D1 - PD_D3 | -3.40 | 0.0245 |
|  |  |  | P_D1 - PD_D3 | -3.32 | 0.0320 |
|  | Ca | χ^2^(8) = 36.64, p = 0, eps2 = 0.872 | P_D1 - P_D3 | -3.88 | 0.0038 |
|  |  |  | P_D1 - P_D3 | -3.50 | 0.0166 |
|  |  |  | P_D1 - PD_D1 | -4.51 | 0.0002 |
|  |  |  | P_D2 - PD_D1 | -3.20 | 0.0497 |
|  |  |  | P_D1 - PD_D1 | -4.10 | 0.0015 |
|  |  |  | P_D2 - PD_D1 | -3.27 | 0.0382 |
|  | Fe | χ^2^(8) = 35.09, p = 0, eps2 = 0.816 | P_D1 - P_D3 | -3.40 | 0.0245 |
|  |  |  | P_D1 - PD_D1 | -3.84 | 0.0044 |
|  |  |  | P_D2 - PD_D1 | -3.50 | 0.0170 |
|  |  |  | P_D1 - PD_D1 | -4.28 | 0.0007 |
|  |  |  | P_D2 - PD_D1 | -3.47 | 0.0187 |
|  | K | χ^2^(8) = 34.3, p = 0, eps2 = 0.837 | P_D1 - P_D3 | -3.48 | 0.0180 |
|  |  |  | P_D2 - P_D3 | -3.81 | 0.0049 |
|  |  |  | P_D1 - P_D3 | -3.72 | 0.0070 |
|  |  |  | P_D2 - P_D3 | -3.30 | 0.0349 |
|  | Mg | χ^2^(8) = 39.02, p = 0, eps2 = 0.929 | P_D1 - P_D3 | -3.68 | 0.0085 |
|  |  |  | P_D1 - P_D3 | -3.55 | 0.0139 |
|  |  |  | P_D1 - PD_D1 | -4.38 | 0.0004 |
|  |  |  | P_D2 - PD_D1 | -3.48 | 0.0184 |
|  |  |  | P_D1 - PD_D1 | -4.21 | 0.0009 |
|  |  |  | P_D2 - PD_D1 | -3.32 | 0.0319 |
|  | P | χ^2^(8) = 26.61, p = 8e-04, eps2 = 0.619 | P_D1 - PD_D2 | 3.99 | 0.0024 |
|  |  |  | P_D2 - PD_D2 | 3.52 | 0.0155 |
|  |  |  | P_D1 - PD_D3 | 3.50 | 0.0170 |
| Sum of fractions | Al | χ^2^(8) = 37.89, p = 0, eps2 = 0.881 | P_D1 - P_D3 | -4.60 | 0.0001 |
|  |  |  | P_D2 - P_D3 | -3.40 | 0.0245 |
|  |  |  | P_D1 - P_D3 | -4.17 | 0.0011 |
|  |  |  | P_D1 - PD_D2 | -3.45 | 0.0204 |
|  |  |  | P_D1 - PD_D3 | -3.74 | 0.0066 |
|  |  |  | P_D1 - PD_D3 | -3.31 | 0.0334 |
|  | Ca | χ^2^(8) = 38.76, p = 0, eps2 = 0.901 | P_D1 - P_D3 | -4.41 | 0.0004 |
|  |  |  | P_D1 - P_D3 | -3.78 | 0.0057 |
|  |  |  | P_D1 - PD_D2 | -4.43 | 0.0003 |
|  |  |  | P_D1 - PD_D2 | -3.80 | 0.0051 |
|  |  |  | P_D1 - PD_D3 | -3.62 | 0.0106 |
|  | Fe | χ^2^(8) = 34.59, p = 0, eps2 = 0.804 | P_D1 - P_D3 | -4.04 | 0.0019 |
|  |  |  | P_D1 - P_D3 | -3.31 | 0.0334 |
|  |  |  | P_D1 - PD_D1 | -3.98 | 0.0025 |
|  |  |  | P_D1 - PD_D3 | -3.25 | 0.0416 |
|  |  |  | P_D1 - PD_D2 | -3.77 | 0.0060 |
|  | K | χ^2^(8) = 39.23, p = 0, eps2 = 0.912 | P_D2 - P_D3 | 3.24 | 0.0433 |
|  |  |  | P_D3 - P_D1 | -3.97 | 0.0026 |
|  |  |  | P_D3 - PD_D1 | -4.39 | 0.0004 |
|  |  |  | P_D3 - PD_D1 | -3.24 | 0.0434 |
|  |  |  | P_D1 - PD_D3 | 3.66 | 0.0092 |
|  |  |  | PD_D1 - PD_D3 | 4.10 | 0.0015 |
|  | Mg | χ^2^(8) = 37.04, p = 0, eps2 = 0.861 | P_D1 - P_D3 | 3.48 | 0.0179 |
|  |  |  | P_D1 - P_D2 | 3.72 | 0.0072 |
|  |  |  | P_D3 - PD_D1 | -3.70 | 0.0077 |
|  |  |  | P_D2 - PD_D1 | -3.95 | 0.0028 |
|  |  |  | P_D1 - PD_D3 | 3.35 | 0.0293 |
|  |  |  | PD_D1 - PD_D3 | 3.58 | 0.0123 |
|  | Zn | χ^2^(8) = 40.64, p = 0, eps2 = 0.945 | P_D1 - P_D3 | -3.46 | 0.0195 |
|  |  |  | P_D1 - PD_D2 | -3.47 | 0.0186 |
|  |  |  | P_D1 - PD_D2 | -4.15 | 0.0012 |
|  |  |  | P_D1 - PD_D3 | -4.09 | 0.0016 |
|  |  |  | P_D1 - PD_D3 | -4.76 | 0.0001 |
|  |  |  | P_D2 - PD_D3 | -3.27 | 0.0381 |
|  | P | χ^2^(8) = 36.07, p = 0, eps2 = 0.839 | P_D1 - P_D3 | 3.79 | 0.0054 |
|  |  |  | P_D2 - P_D3 | 3.23 | 0.0453 |
|  |  |  | P_D1 - PD_D2 | 3.84 | 0.0044 |
|  |  |  | P_D2 - PD_D2 | 3.27 | 0.0381 |
|  |  |  | P_D1 - PD_D3 | 3.82 | 0.0049 |
|  |  |  | P_D2 - PD_D3 | 3.25 | 0.0416 |

### Table S8- Significant (p < 0.05) Kruskal-Wallis and post-hoc Dunn’s test results for pairwise comparisons of mean soil exoenzyme activity between treatment and timepoint groupings. Groups used for comparisons include the no dust control at T1 or T2 months (P_T1; P_T2), and the dust treatment at T1 or T2 months (PD_T1; PD_T2). Test results are organized by the type of exoenzyme.

|  |  |  | Dunn’s post hoc result | |
| --- | --- | --- | --- | --- |
| Enzyme | KW | Comparison | Z statistic | p-value^BF^ |
| Glucosidase | χ^2^(3) = 27.19, p = 0, ε^2^ = 0.367 | P_T1 - P_T2 | 2.96 | 0.0182 |
|  |  | P_T1 - PD_T1 | 4.47 | 0.0000 |
|  |  | PD_T1 - PD_T2 | -4.07 | 0.0003 |
| Phosphatase | χ^2^(3) = 40.66, p = 0, ε^2^ = 0.549 | P_T1 - PD_T2 | 5.81 | 0.0000 |
|  |  | P_T2 - PD_T2 | 3.18 | 0.0089 |
|  |  | PD_T1 - PD_T2 | 5.12 | 0.0000 |
| Xylosidase | χ^2^(3) = 9.14, p = 0.0275, ε^2^ = 0.124 | PD_T1 - PD_T2 | -2.72 | 0.0394 |

### Table S9- Significant (p < 0.05) Kruskal-Wallis and post-hoc Dunn’s test results for pairwise comparisons of mean soil exoenzyme activity between unique treatment-depth groupings. Treatments included the control at T0 (P), no dust control at T2 months (P), and the dust treatment at T2 months (PD_T2). Depths included the upper (D1), middle (D2), and lower (D3) peat layers. Test results are organized by the type of exoenzyme.

|  |  |  | Dunn’s post hoc result | |
| --- | --- | --- | --- | --- |
| Enzyme | KW | Comparison | Z statistic | BF corrected p-value |
| Glucosidase | χ^2^(8) = 32.36, p = 1e-04, ε^2^ = 0.437 | P_D3 - P_D1 | 3.30 | 0.0347 |
|  |  | P_D3 - P_D3 | 3.49 | 0.0172 |
|  |  | P_D2 - PD_D1 | 3.45 | 0.0204 |
|  |  | P_D3 - PD_D1 | 4.21 | 0.0009 |
|  |  | P_D3 - PD_D3 | 3.67 | 0.0088 |
| NAGase | χ^2^(8) = 50.12, p = 0, ε^2^ = 0.677 | P_D1 - P_D2 | 4.24 | 0.0008 |
|  |  | P_D2 - P_D2 | 4.37 | 0.0004 |
|  |  | P_D3 - P_D2 | 4.27 | 0.0007 |
|  |  | P_D1 - PD_D1 | 3.72 | 0.0073 |
|  |  | P_D2 - PD_D1 | 3.85 | 0.0043 |
|  |  | P_D3 - PD_D1 | 3.75 | 0.0064 |
|  |  | P_D1 - PD_D3 | 3.95 | 0.0029 |
|  |  | P_D2 - PD_D3 | 4.08 | 0.0016 |
|  |  | P_D3 - PD_D3 | 3.98 | 0.0025 |
| Phosphatase | χ^2^(8) = 30.71, p = 2e-04, ε^2^ = 0.415 | P_D1 - PD_D1 | 4.31 | 0.0006 |
|  |  | P_D3 - PD_D1 | 3.90 | 0.0034 |
|  |  | P_D3 - PD_D1 | 3.68 | 0.0083 |
| Xylosidase | χ^2^(8) = 42.57, p = 0, ε^2^ = 0.575 | P_D1 - P_D2 | -3.51 | 0.0164 |
|  |  | P_D1 - P_D3 | -3.46 | 0.0191 |
|  |  | P_D2 - PD_D1 | 3.60 | 0.0114 |
|  |  | P_D3 - PD_D1 | 3.55 | 0.0139 |
|  |  | P_D1 - PD_D2 | -3.61 | 0.0110 |
|  |  | PD_D1 -  PD_D2 | -3.73 | 0.0069 |
|  |  | P_D1 - PD_D3 | -3.34 | 0.0299 |
|  |  | PD_D1 –  PD_D3 | -3.40 | 0.0241 |

### Table S10- Significant (p < 0.05) Kruskal-Wallis and post-hoc Dunn’s test results for pairwise comparisons of mean absorbance on Biolog Ecoplates^TM^ between treatment and timepoint groupings. Groups used for comparisons include the no dust control at T1 or T2 months (P_T1; P_T2), and the dust treatment at T1 or T2 months (PD_T1; PD_T2). Test results are organized by the functional group of the carbon substrates in the Ecoplates^TM^.

|  |  |  | Dunn’s post hoc result | |
| --- | --- | --- | --- | --- |
| Functional Group | KW | Comparison | Z statistic | p-value^BF^ |
| Amines | χ^2^(4) = 29.54, p = 0, ε^2^ = 0.671 | P_T0 - P_T1 | 2.99 | 0.0278 |
|  |  | P_T0 - PD_T1 | 3.73 | 0.0019 |
|  |  | P_T2 - PD_T1 | 3.49 | 0.0047 |
|  |  | P_T1 - PD_T2 | -3.48 | 0.0051 |
|  |  | PD_T1 - PD_T2 | -4.21 | 0.0003 |
| Amino Acids | χ^2^(4) = 32.34, p = 0, ε^2^ = 0.735 | P_T0 - P_T1 | 3.22 | 0.0127 |
|  |  | P_T1 - P_T2 | -3.31 | 0.0093 |
|  |  | P_T0 - PD_T1 | 3.55 | 0.0039 |
|  |  | P_T2 - PD_T1 | 3.64 | 0.0028 |
|  |  | P_T1 - PD_T2 | -3.89 | 0.0010 |
|  |  | PD_T1 - PD_T2 | -4.21 | 0.0003 |
| Carbohydrates | χ^2^(4) = 34.6, p = 0, ε^2^ = 0.786 | P_T1 - P_T2 | -3.46 | 0.0053 |
|  |  | P_T0 - PD_T1 | 3.25 | 0.0116 |
|  |  | P_T2 - PD_T1 | 4.34 | 0.0001 |
|  |  | P_T1 - PD_T2 | -3.75 | 0.0018 |
|  |  | PD_T1 - PD_T2 | -4.63 | 0.0000 |
| Carboxylic Acids | χ^2^(4) = 34.93, p = 0, ε^2^ = 0.794 | P_T0 - P_T1 | 4.43 | 0.0001 |
|  |  | P_T0 - PD_T1 | 4.97 | 0.0000 |
|  |  | P_T2 - PD_T1 | 3.28 | 0.0102 |
|  |  | PD_T1 - PD_T2 | -3.32 | 0.0090 |
| Phenolics | χ^2^(4) = 29.58, p = 0, ε^2^ = 0.672 | P_T1 - P_T2 | -3.94 | 0.0008 |
|  |  | P_T2 - PD_T1 | 4.24 | 0.0002 |
|  |  | P_T1 - PD_T2 | -3.36 | 0.0077 |
|  |  | PD_T1 - PD_T2 | -3.67 | 0.0024 |
| Polymers | χ^2^(4) = 32.88, p = 0, ε^2^ = 0.747 | P_T0 - P_T1 | 3.37 | 0.0074 |
|  |  | P_T0 - PD_T1 | 4.70 | 0.0000 |
|  |  | P_T1 - PD_T2 | -3.18 | 0.0149 |
|  |  | PD_T1 - PD_T2 | -4.51 | 0.0001 |
| TOTAL | χ^2^(4) = 33.06, p = 0, ε^2^ = 0.751 | P_T0 - P_T1 | 3.34 | 0.0084 |
|  |  | P_T1 - P_T2 | -2.84 | 0.0457 |
|  |  | P_T0 - PD_T1 | 3.93 | 0.0008 |
|  |  | P_T2 - PD_T1 | 3.43 | 0.0061 |
|  |  | P_T1 - PD_T2 | -3.84 | 0.0012 |
|  |  | PD_T1 - PD_T2 | -4.43 | 0.0001 |

### Table S11- Diversity measurements for prokaryotic communities throughout the 2-month peat-mesocosm experiment. Each sample is representative of 5 extraction replicates pooled together (prior to sequencing). Samples are organized by timepoint (T0, T1, and T2 months), peat depth (upper (D1), middle (D2), lower (D3)), and treatment (peat without dust (P) and peat with dust (PD)).

| Time | Depth | Treatment | Richness | Berger-Parker Index | Shannon Diversity Index |
| --- | --- | --- | --- | --- | --- |
| T0 | D1 | P | 1159 | 0.035422 | 6.303237 |
|  | D2 |  | 863 | 0.014772 | 6.262344 |
|  | D3 |  | 1127 | 0.026864 | 6.366374 |
| T1 | D1 | P | 986 | 0.030522 | 6.301948 |
|  |  | PD | 868 | 0.01425 | 6.218617 |
|  | D2 | P | 737 | 0.01929 | 6.130936 |
|  |  | PD | 980 | 0.025454 | 6.385361 |
|  | D3 | P | 1099 | 0.017945 | 6.430176 |
|  |  | PD | 921 | 0.032934 | 6.183489 |
| T2 | D1 | P | 650 | 0.014951 | 6.024025 |
|  |  | PD | 478 | 0.03184 | 5.673563 |
|  | D2 | P | 958 | 0.015125 | 6.328239 |
|  |  | PD | 862 | 0.012104 | 6.266199 |
|  | D3 | P | 757 | 0.021625 | 6.062468 |
|  |  | PD | 748 | 0.018719 | 6.154683 |

### Table S12- Diversity measurements for fungal communities throughout the 2-month peat-mesocosm experiment. Each sample is representative of 5 extraction replicates pooled together (prior to sequencing). Samples are organized by timepoint (T0, T1, and T2 months), peat depth (upper (D1), middle (D2), lower (D3)), and treatment (peat without dust (P) and peat with dust (PD)). Due to a low sequencing depth, diversity metrics are not reported for the PD sample from D2 at T1 month.

| Time | Depth | Treatment | Richness | Berger-Parker Index | Shannon Diversity Index |
| --- | --- | --- | --- | --- | --- |
| T0 | D1 | P | 428 | 0.196267 | 2.843824 |
|  | D2 |  | 302 | 0.217203 | 2.545245 |
|  | D3 |  | 549 | 0.277117 | 3.213794 |
| T1 | D1 | P | 321 | 0.30512 | 3.123562 |
|  |  | PD | 364 | 0.162754 | 3.173097 |
|  | D2 | P | NA | NA | NA |
|  |  | PD | 131 | 0.363334 | 2.757514 |
|  | D3 | P | 323 | 0.157738 | 3.207387 |
|  |  | PD | 324 | 0.135471 | 3.497972 |
| T2 | D1 | P | 422 | 0.126701 | 3.426967 |
|  |  | PD | 352 | 0.117235 | 3.816594 |
|  | D2 | P | 560 | 0.15552 | 3.61433 |
|  |  | PD | 356 | 0.129969 | 3.28024 |
|  | D3 | P | 388 | 0.238712 | 3.207348 |
|  |  | PD | 437 | 0.154305 | 3.557838 |

### Table S13- Beta-dispersion (BETADISP) test results for fungal and bacterial communities at the ASV level (n permutations = 999). These results were consistent when count data was agglomerated to the genus or family taxonomic rank. Significant results (p < 0.05) are bolded and have an asterisk (*).

| Microbial group | Taxonomic Level | Parameter | Df | SumOfSqs | Mean Sq | F | p-value |
| --- | --- | --- | --- | --- | --- | --- | --- |
| Bacteria (16S) | Family | Treatment | 1 | 57 | 57 | 2.10 | 0.149 |
|  |  | Time | 2 | 20 | 10 | 0.23 | 0.822 |
|  |  | Depth | 2 | 139 | 69 | 1.91 | 0.194 |
| Fungi (ITS gene) | Genus | Treatment | 1 | 13 | 13 | 0.38 | 0.553 |
|  |  | **Time** | **2** | **726** | **363** | **12.59** | **0.003*** |
|  |  | Depth | 2 | 117 | 59 | 1.74 | 0.234 |

### Table S14- The relative abundances (%) of each trophic mode in peat samples. Calculated abundances are relative to the total community in a given sample, regardless if there was a match in the FUNGuild dataset or not. Guilds were assigned with the FUNGUILD database (version 1.1) and plotted with the *microeco* R package. Saprotroph-Symbiotroph-Pathotroph

| Time | Depth | Treatment | Saprotroph | Symbiotroph | Pathotroph | Saprotroph-Symbiotroph | Saprotroph-Pathotroph | Symbiotroph-Pathotroph | Saprotroph-Symbiotroph-Pathotroph |
| --- | --- | --- | --- | --- | --- | --- | --- | --- | --- |
| T1 | D1 | PD | 35.2 | 23.2 | 0.05 | 1.59 | 0.04 | 0.36 | 2.90 |
|  |  | P | 16.7 | 18.5 | 0.67 | 3.00 | 0.41 | 0.47 | 0.61 |
|  | D2 | PD | 50.9 | 20.9 | 0.16 | 1.04 | 0.03 | 0.14 | 0.85 |
|  |  | P | 28.1 | 11.9 | 0.10 | 4.93 | 0.23 | 0.06 | 2.04 |
|  | D3 | PD | 22.9 | 39.0 | 0.39 | 2.41 | 0.18 | 0.33 | 1.05 |
|  |  | P | 34.5 | 6.7 | 0.07 | 1.90 | 0.08 | 0 | 4.15 |
| T2 | D1 | PD | 36.3 | 19.0 | 0.08 | 0.44 | 0.67 | 0.41 | 4.75 |
|  |  | P | 27.6 | 17.1 | 0.05 | 3.59 | 0.76 | 0.26 | 4.71 |
|  | D2 | PD | 38.4 | 15.0 | 0.04 | 1.87 | 0.05 | 0.33 | 1.13 |
|  |  | P | 12.4 | 9.8 | 0.21 | 4.32 | 0.25 | 0.09 | 14.92 |
|  | D3 | PD | 28.2 | 21.7 | 0.67 | 12.44 | 0.43 | 0.03 | 2.32 |
|  |  | P | 31.7 | 9.9 | 0.12 | 3.08 | 0.28 | 0.02 | 0.96 |
