## Supplemental Figures for "Mineral dust stimulates microbial exoenzyme activity and enhances carbon mineralization capabilities in nutrient-poor peat soil"

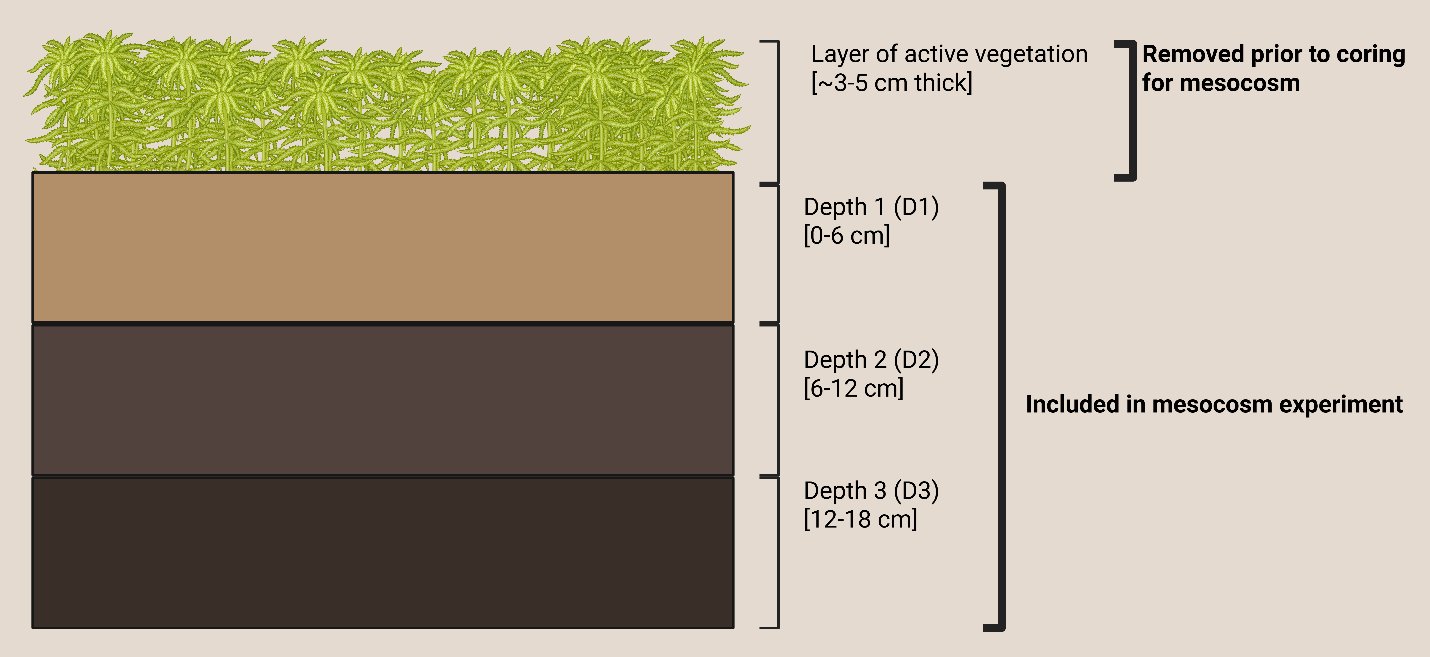

Figure S1- Simple illustration of peat layers included or removed prior to constructing the mesocosm experiments. This figure was made with BioRender.com.

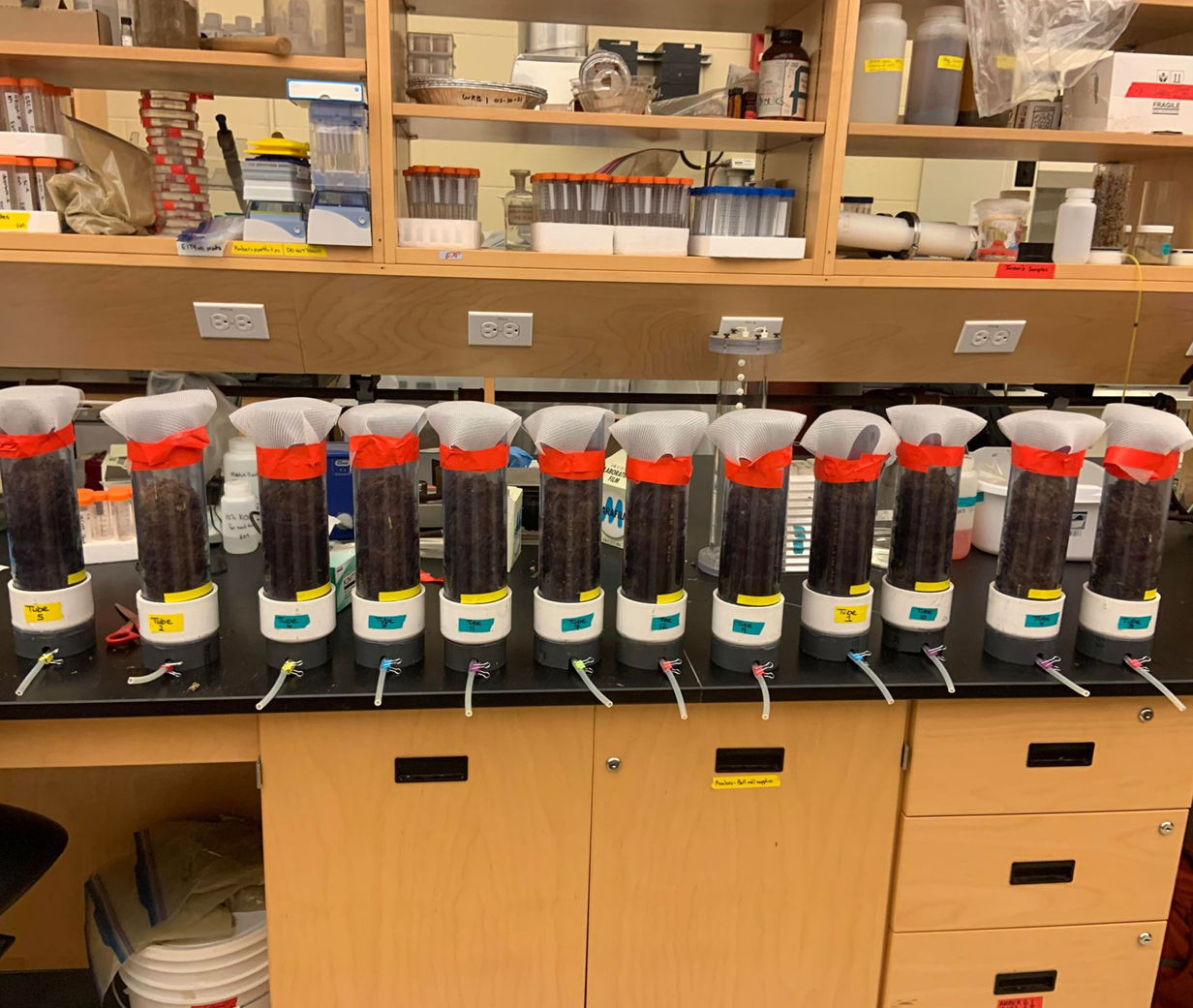

Figure S2- Peat mesocosms prior to incubation. Each mesocosm core was covered at the top with nylon mesh and capped at the bottom with an outlet for porewater drainage.

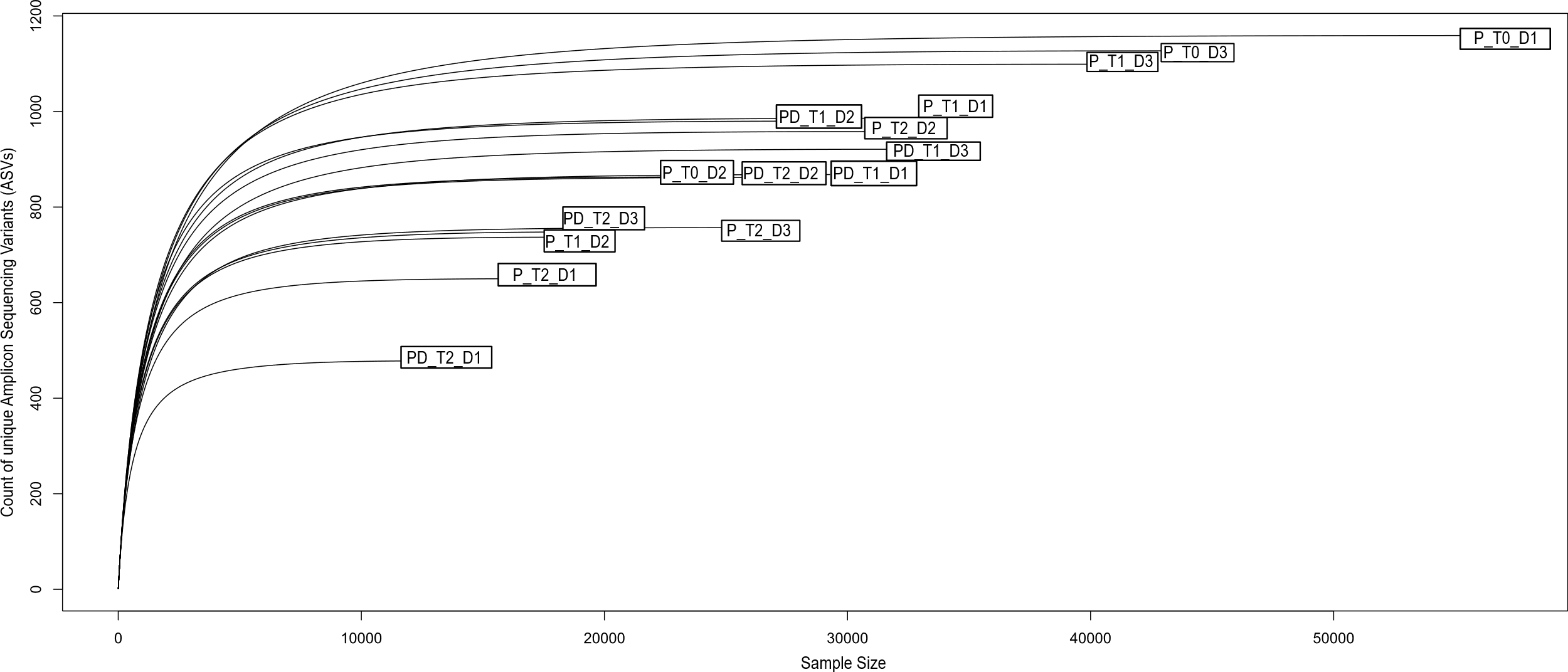

Figure S3- Rarefaction curve for 16S rRNA gene amplicons from peat samples. Sample identifiers indicate the treatment (peat with no dust [P] and peat with dust [PD]), the timepoint (T0, T1, and T2 months), and the peat depth from the top of the core (D1: 0-6, D2: 6-12, and D3: 12-18 cm).

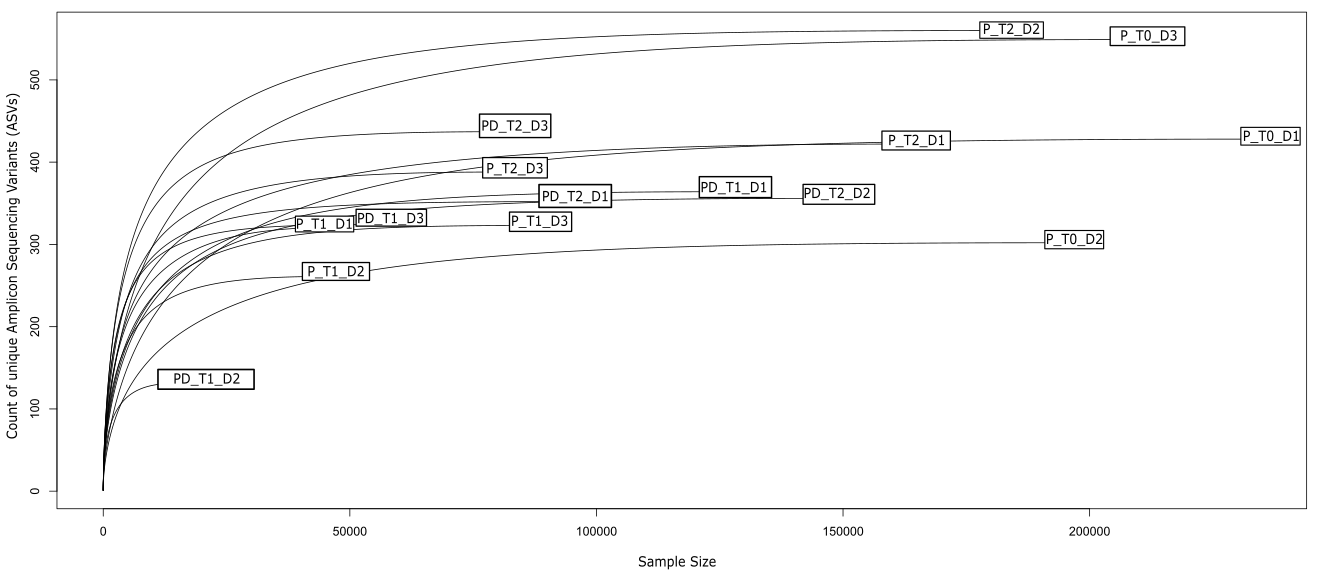

Figure S4- Rarefaction curve for ITS gene amplicons from peat samples. Sample identifiers indicate the treatment (peat with no dust [P] and peat with dust [PD]), the timepoint (T0, T1, and T2 months), and the peat depth from the top of the core (D1: 0-6, D2: 6-12, and D3: 12-18 cm).

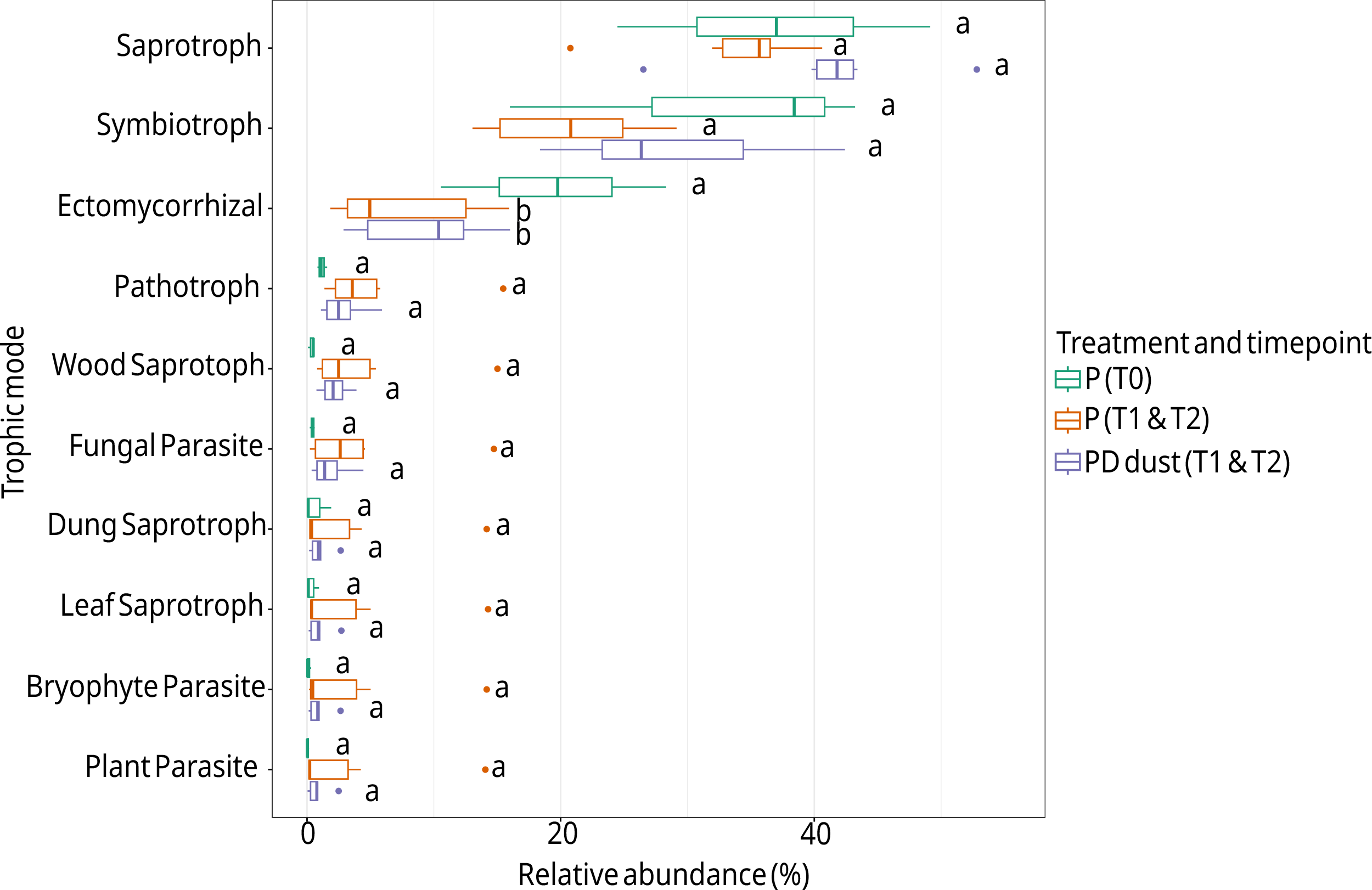

Figure S5- Average relative abundance of fungal trophic modes between treatments. Trophic modes were predicted with the FUNGuild database (version 1.1). For a given treatment (peat with no dust [P] and peat with dust [PD]), relative abundances were averaged across all depths and timepoints (T1 month and T2 months). Values with different letters indicate statistically significant differences in the relative abundance of a given trophic mode (Duncan’s post hoc test).
